# Mutational basis of serum cross-neutralization profiles elicited by infection or vaccination with SARS-CoV-2 variants

**DOI:** 10.1101/2023.08.13.553144

**Authors:** Kshitij Wagh, Xiaoying Shen, James Theiler, Bethany Girard, Jean-Claude Marshall, David C. Montefiori, Bette Korber

## Abstract

A series of SARS-CoV-2 variants emerged during the pandemic under selection for neutralization resistance. Convalescent and vaccinated sera show consistently different cross-neutralization profiles depending on infecting or vaccine variants. To understand the basis of this heterogeneity, we modeled serum cross-neutralization titers for 165 sera after infection or vaccination with historically prominent lineages tested against 18 variant pseudoviruses. Cross-neutralization profiles were well captured by models incorporating autologous neutralizing titers and combinations of specific shared and differing mutations between the infecting/vaccine variants and pseudoviruses. Infecting/vaccine variant-specific models identified mutations that significantly impacted cross-neutralization and quantified their relative contributions. Unified models that explained cross-neutralization profiles across all infecting and vaccine variants provided accurate predictions of holdout neutralization data comprising untested variants as infecting or vaccine variants, and as test pseudoviruses. Finally, comparative modeling of 2-dose versus 3-dose mRNA-1273 vaccine data revealed that the third dose overcame key resistance mutations to improve neutralization breadth.

**HIGHLIGHTS:**

- Modeled SARS-CoV-2 cross-neutralization using mutations at key sites
- Identified resistance mutations and quantified relative impact
- Accurately predicted holdout variant and convalescent/vaccine sera neutralization
- Showed that the third dose of mRNA-1273 vaccination overcomes resistance mutations

## INTRODUCTION

In the more than three years of the COVID-19 pandemic, continued viral evolution subjected to the forces of natural selection for improved transmission and immune escape has led to a series of major global transitions between variant lineages (Fig. S1). These were historically designated by the World Health Organization (WHO) as “variants of concern” (VOCs), “variants of interest” (VOI) or “variants under monitoring” (VUMs), using Greek letter names for dominant lineages from Alpha through Omicron^1^. Omicron has continued to evolve into many sublineages; currently the Pango lineages^2^ referred to as XBB.1.5 and XBB.1.16 are the most globally dominant forms, and considered to be VOIs, with several additional Omicron subvariants classified as VUMs^1^. These VOCs and VOIs have accounted for a substantial fraction of the total toll on human health and death, with their introductions historically often associated with “waves” of increased infections (Fig. S1)^3^.

Previous and emerging variants have a disproportionately high number of mutations in the *spike* gene^4^ indicative of selection pressures resulting from neutralizing antibody escape^5–11^, improved ACE2 binding^12,13^, increased Spike cleavage^14^, resistance to innate immune responses^15^, or a combination thereof^16^. Neutralizing antibody (NAb) responses correlate with protection from SARS-CoV-2 infection and disease^17–19^, but recent and emergent variants continue to evolve increasing resistance to such responses^10,20,21^. Still, immunity conferred by prior infection and vaccination is beneficial; in breakthrough cases (i.e. infections in vaccinated individuals), immune responses continue to help protect against progression to severe disease^22^ and reduce transmission risk^23^. Taken together, this suggests that the population-level immunity derived from past vaccination and infection history can modulate the dominance of a particular variant in that population, in addition to the number and nature of co-circulating variants and epidemiological factors^3^. At the time of writing, the pandemic is at a complex stage of co-circulation of multiple Omicron sublineage variants that are referred to by their Pango lineage designations^2^ (Fig. S1). The recombinant Omicron sublineage XBB.1.5 is currently the predominant variant in much of the world and continues to be increasingly sampled. However, newer variants (e.g., XBB.1.16, XBB.1.9, XBB.2.3, and XBC.1.6…) are each increasing at a faster pace than XBB.1.5, indicative of a relative transmission advantage (Fig. S1). All XBB lineages carry an array of mutations that confer resistance to neutralization by sera from infections with past variants and vaccination^12,24,25^, but XBB.1.5 and other expanding variants also carry the Spike S486P mutation that by enhancing binding to the ACE2 receptor may compensate for a fitness loss from other mutations^12,24^. The selective advantage of XBB.1.5 illustrates that SARS-CoV-2 continues to adapt to human hosts under the same selective forces that have dictated variant transitions in the past.

Many studies have tracked the sensitivity of variants to neutralization by vaccine-induced responses or monoclonal antibodies^5–9,26,27^. Others have focused on the isolation of monoclonal antibodies that target epitopes enabling broad cross-recognition even beyond SARS-CoV-2^28–33^. Lesser information is available on the neutralizing responses induced by infection by such variants^34,35^, and on the systematic comparison of neutralization profiles induced by infection by different variants^36,37^. These studies have documented the consistent differences in cross-neutralization profiles of convalescent sera post-infection with different variants and post-vaccination, indicating that different variants induce distinctive neutralizing responses. While these studies have elucidated relationships between the infecting/vaccine variants in terms of the neutralization responses they induce, they did not provide direct information about the nature of the epitopes targeted in the context of different variants. Deep mutational scanning (DMS) ^34,35,38,39^ and other neutralization mapping studies^10,21^ have provided information about resistance mutations to neutralization responses by convalescent or vaccinated sera, but these studies did not quantitatively evaluate the relative contribution to resistance profiles resulting from mutations that often recur in combination across multiple VOCs.

Variant-specific differences in cross-neutralization profiles could arise from the induced neutralizing antibodies (NAbs) targeting variant-specific amino acids/glycans within a given epitope, or shifts in epitope accessibility or immunodominance^16,34^ across variants. In this work, we hypothesized that key aspects of these scenarios could be captured by comparing the constellation of mutations in the infecting or vaccine variants that elicit the response and the pseudovirus variants used for testing neutralization breadth. We investigated this hypothesis using an extensive cross-neutralization dataset for two-dose mRNA-1273 vaccinated sera and convalescent sera post-primary infection with SARS-CoV-2 by one of 7 major variants, tested against 18 pseudoviruses encompassing 11 major variants. By applying statistical modeling to these data we showed that cross-neutralization for sera from each infecting or vaccine variant could be accurately modeled as a function of amino acid patterns at key sites, when combined with neutralizing titers of the infecting variant-matched pseudovirus (“autologous titers”) that serve as a proxy for the strength of neutralizing responses from each participant. Next, we showed that a model that encompassed mutations in both infecting/vaccine and pseudovirus variants well captured cross-neutralization profiles across all infecting and vaccine variants and accurately predicted holdout data (i.e. data that model was not trained on) from novel vaccine/infecting and pseudovirus variants. Finally, a comparison of 2-dose versus 3-dose mRNA-1273 vaccine data showed that the third dose enabled overcoming key resistance mutations leading to improved neutralization breadth.

## RESULTS

### Serum neutralization data shows infecting/vaccine variant-specific heterogeneous cross-neutralization profiles

We collected serum neutralization data for 165 sera post-vaccination with mRNA-1273 (2 doses), or post-primary infection with D614G, or one of six historical variants of concern or interest (Alpha, Beta, Gamma, Delta, Iota, and Lambda). The cohort and geographic location information for these samples is presented in Table S1. These sera were tested against a total of 18 pseudoviruses that included D614G; Alpha and Alpha+E484K (here “E484K” indicates that “E” (glutamic acid) is the amino acid found in in the ancestral, 484 is the position in Spike, and “K” (lysine) is the mutated amino acid and the “+” indicates that this mutation was not present in the baseline Alpha sequence); Beta; Gamma; five Delta variants, AY.1, AY.2, AY.3, AY.3+E484Q and AY.3+K417N; Iota; Lambda; Epsilon; Kappa; Mu; Omicron BA.1 and BA.1+A_BA.1_484K (where the non-ancestral E484A (A_BA.1_) in the baseline BA.1 sequence was replaced by E484K); and Omicron BA.2 (Fig. 1A, Table S2). The defining Spike mutations found in each baseline variant are included in Fig. S2. Although sample availability prohibited testing each serum against all the pseudoviruses, each serum was tested against a median of 15 pseudoviruses. The sera showed a spectrum of cross-neutralization breadth and potency, with the highest neutralizing titers (measured as 50% inhibitory dilution or ID_50_) generally found against the autologous pseudovirus (i.e., matching the infecting or vaccine variant) (Fig. 1B). Both breadth and potency of cross-neutralization for each serum were significantly and positively correlated with autologous ID_50_ (Fig. 1B-C), indicating that autologous neutralization potency serves as a significant, albeit limited, baseline surrogate of cross-neutralization potential, and that potency can overcome some of the barriers to breadth.

**Figure 1:**
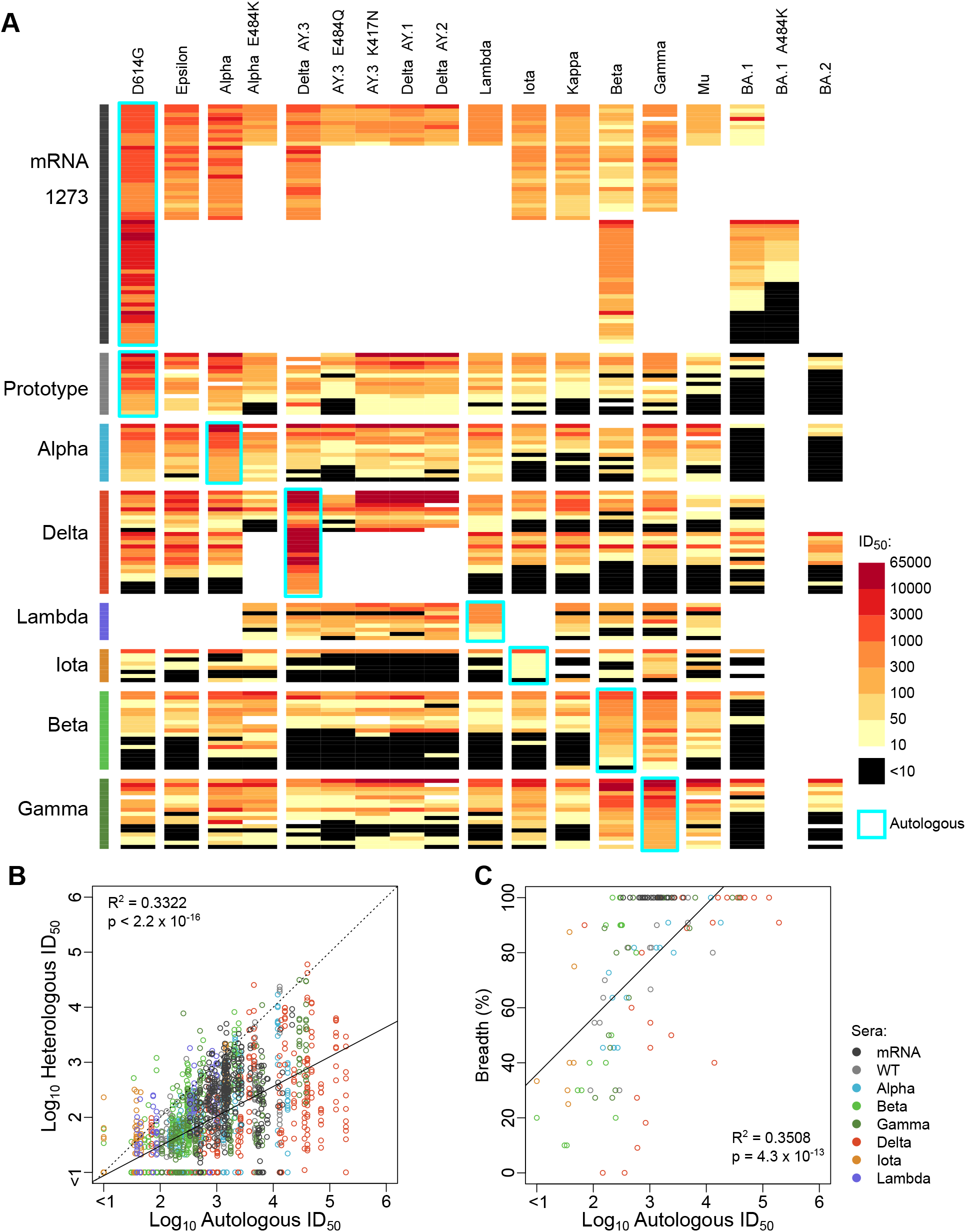
Serum neutralization data. (A) A heatmap displaying neutralizing ID_50_ titers, with sera grouped according to their vaccine or infecting variant as rows and pseudovirus variants as columns. Autologous titers are indicated in cyan boxes. Black cells indicate ID50 values below threshold (<10) and blank cells indicate no data. (B) Heterologous ID50 titers (vertical axis) for each serum are plotted as a function of autologous ID_50_ titers (horizontal axis) color coded per the legend in (C). Dotted line indicates identity and solid line indicates linear fit. Coefficient of determination, R^2^, and p-value from Kendall tau rank test are shown. (C) Same as (B), except breadth of cross-neutralization (fraction of tested viruses with ID_50_ > 10) is shown on the vertical axis.

Sera from the same infecting/vaccine variant exhibited similar cross-neutralization patterns, but sera from different infecting/vaccine variants were distinctive (Fig. 2A). For example, sera from Gamma and Iota infections cross-neutralized Beta with comparable titers to the autologous variant, while most other sera showed significantly reduced titers against Beta, such as the Delta-infection sera, most of which had titers below threshold (ID_50_ < 10) against the Beta pseudovirus (Fig. 2A). The cross-neutralization propensity between a given infecting/vaccine variant and a given pseudovirus variant was roughly symmetric. For example, most Beta-infection sera also showed below threshold titers (ID_50_ < 10) against the Delta pseudovirus, mirroring the resistance of Beta pseudovirus to Delta-infection sera.

**Figure 2:**
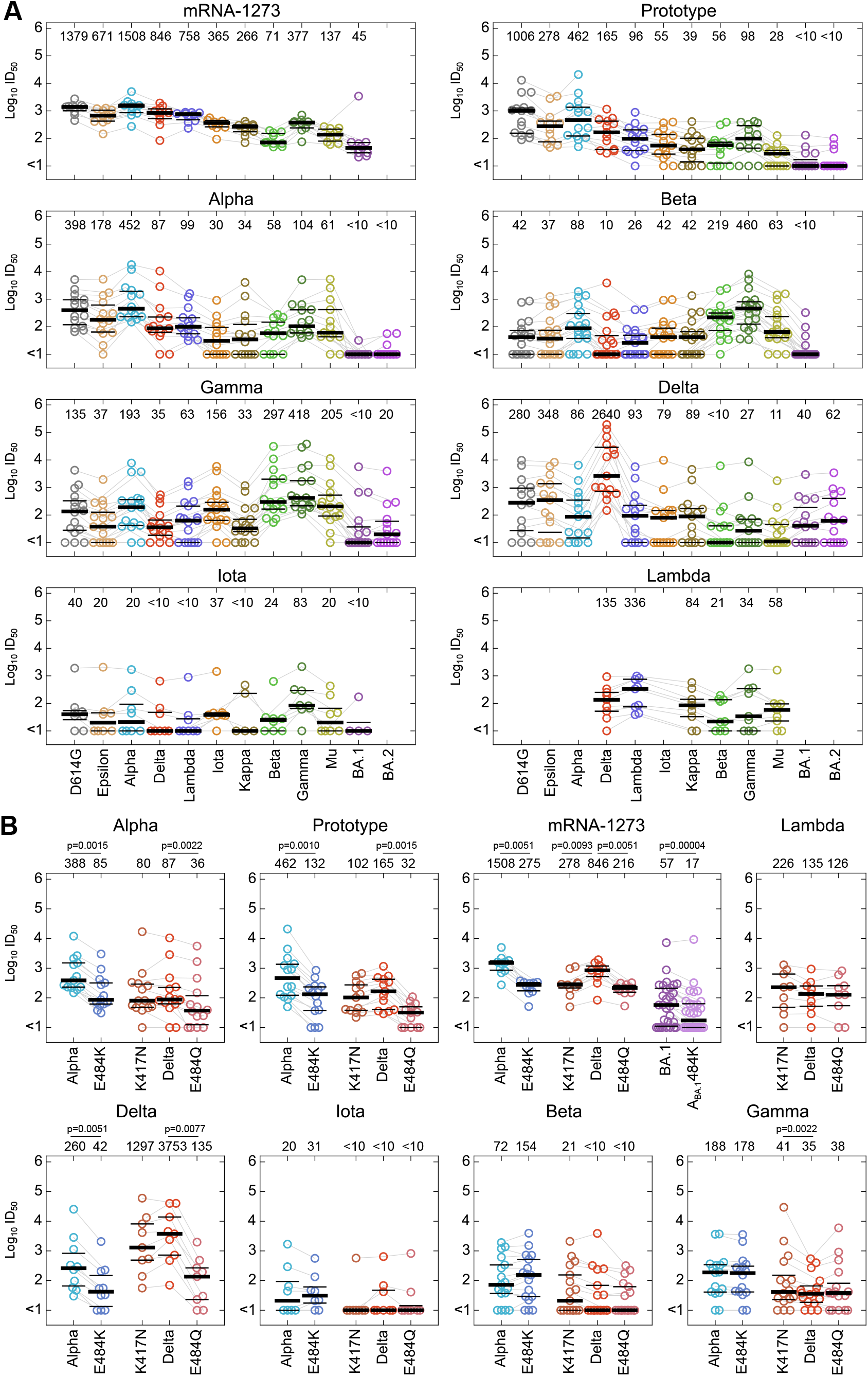
(A) Subpanels show pooled ID_50_ titers for sera from each infecting/vaccine variant group against the pseudoviruses tested. Median titers are noted on the top of each sub-panel and medians (thick horizontal lines) and first and third quartiles (thin horizontal lines) are included. Light grey lines connect the data points from the same serum between consecutive pseudoviruses. (B) Same as (A) except with titers against wildtype and mutant pseudoviruses. The latter are labeled by the mutation that was introduced (e.g. E484K for Alpha+E484K) following the convention of ancestral amino acid, position in Spike and mutation, with the exception of A_BA.1_484K that refers to the non-ancestral A484 amino acid in BA.1 mutated to K484. Statistically significant for differences were calculated using two-sided Wilcoxon paired rank test; those with an uncorrected p < 0.05 are indicated.

Pseudovirus point-mutants representing recurrent within-lineage mutations allowed direct quantification of the dependence of serum cross-neutralization on mutations at sites 417 and 484 (Fig. 2B). The K417N in a Delta Spike backbone significantly impacted potency in two cases: mRNA-1273 vaccinated sera showed a 4-fold reduction, while sera from infections with the Gamma variant (which contains the K417T mutation away from ancestral (Fig. S2)) showed a 1.2-fold improvement. K417N is the most common form among prevalent Omicron variants.

Mutations at site 484 had greater impact (Fig. 2B). Median neutralization titers of Prototype (i.e. D614G variants prior to the emergence of Alpha), Alpha, and Delta infection and mRNA-1273 vaccine sera were reduced 3.5-8.6-fold when E484K was introduced in an Alpha backbone, and E484Q in a Delta backbone reduced titers 2.4-3.9-fold for Alpha, Prototype and mRNA-1273 sera, and 27-fold for Delta sera; all these differences were significant. The impact of the A_BA.1_484K mutation in the BA.1 pseudovirus backbone was only tested for mRNA-1273 sera, and a significant 3.3-fold drop was observed. Lambda was the only ancestral E484 carrying infecting variant in our dataset for which serum cross-neutralization was not impacted by 484 mutations. In contrast, responses to infection by Beta or Iota variants that carry the non-ancestral E484K mutation showed modest but non-significant 1.5-2.1-fold increase in titers when E484K was matched in the Alpha pseudovirus background, and responses to infection by Gamma, which also carried E484K, showed no increase. Sera from infections with these E484K-carrying variants were also not impacted by the introduction of E484Q in Delta pseudoviruses. The relevance of mutations at site 484 is discussed in more detail below, but these results suggest that with the exception of Lambda, infecting/vaccine variants with the ancestral E484 could lead to immunodominant responses targeting the 484 site, while infecting variants with E484K may preferentially induce antibodies targeting different epitopes, as suggested previously^34^. Thus, the E484K mutation in the infecting/vaccine variants may contribute to the distinctive cross-neutralization profiles induced by SARS-CoV-2 variants. Omicron viruses have further transitioned to E484A that has been the basic form found in all key Omicron subvariants.

### Mutational variables for neutralization modeling

We hypothesized that shared and non-shared mutations at specific Spike sites between the infecting and test variants may underlie the heterogeneous infecting or vaccine-variant dependent cross-neutralization profiles. To evaluate this hypothesis, we first selected Spike sites that have mutations across multiple variants to include in our models. Our dataset included 18 variants; therefore, it can support a theoretical maximum of 17 linearly independent variables (plus one constant), far fewer than the total number of distinct Spike mutations in the viruses in our dataset (Fig. S2A). Thus, we downselected sites for modeling (Fig. 3A, S2B) using the strategy detailed in STAR methods. Briefly, we selected sites that were mutated in at least 3 variants. Some of these sites showed mutational patterns that co-occurred exactly with mutations at other site(s) in our dataset; e.g., the deletion at sites 69-70 always co-occurred with a deletion at either site 142 alone or at sites 142-144 (Fig. S2C). Such redundancies were excluded, leaving 9 chosen sites with a total of 16 mutations in our dataset (Fig. 3A). Four sites had a single mutation away from the ancestral form (T95I, G142D, T478K, N501Y), 3 sites had 2 mutations (S371L/F, K417T/N, L452R/Q) and 2 sites had 3 mutations (Y144S/del144/del142-144 and E484K/Q/A). Mutations in each of these sites have been shown to impact neutralization by monoclonal NAbs^10,21,40–42^.

**Figure 3:**
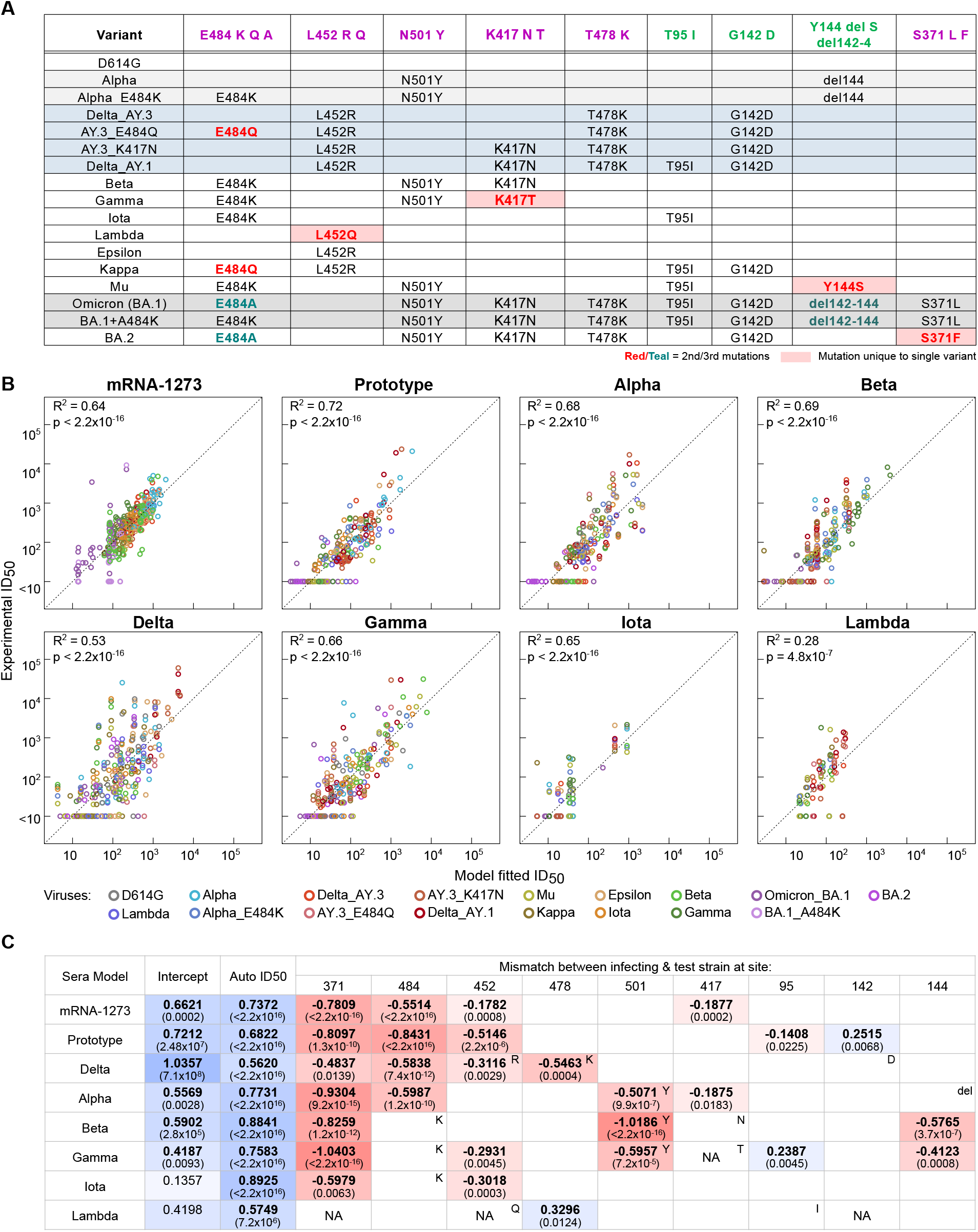
Infecting/vaccine variant specific models. (A) Variant sequences at Spike sites chosen for models. (B) Model fit ID50 titers (horizontal axis) are plotted with experimental ID_50_ titers (vertical axis) for each infecting/vaccine variant model, with colors indicating pseudoviruses. Coefficient of determination (R^2^) and two-sided p-value from Kendall Tau rank test are shown. (C) Coefficients of the best-fit models for each variable (columns) are shown for each infecting/vaccine variant model (rows). Red, white, and blue color scale indicates negative, 0, and positive values, respectively. Coefficients that were significantly different from 0 (p < 0.05, Student’s t-test) are shown in bold and p-values are indicated in parentheses. Blank cells indicate variables dropped in BIC model selection, and ‘NA’ indicates variables with no variation or that are redundant with other variables in the subset of data for a given infecting/vaccine variant. Infecting variant mutations at each site, if different from ancestral, are shown in top right of each cell.

### Mutational differences in infecting and test variants can explain serum cross-neutralization profiles from each infecting/vaccine variant

We next modeled Log_10_ heterologous (i.e., cross-neutralizing) ID_50_ titers for sera from each infecting/vaccine variant as a linear function of Log_10_ autologous ID_50_, which was a significant correlate of heterologous ID_50_ titers (Fig. 1B), and of the sequence variables encoding mutations at the 9 chosen sites. We tested two classes of models that differed in the encoding of mutational variables: i) binary encoding of each mutation at each site in the pseudovirus variant (“pseudovirus mutation model”), and ii) a binary variable indicating mismatch between infecting and pseudovirus variant mutations at each site (“mutation mismatch model”). The former captures the potential impact of exact mutations in the variant pseudoviruses on cross-neutralization, while the latter captures the impact of mutational differences in pseudoviruses away from the infecting/vaccine variant.

To identify the strongest contributing sequence variables, we performed stepwise model reduction using Bayesian Information Criterion (BIC) that removes variables that do not improve model fits above a threshold^43^. For each set of models, the BIC-reduced models were not significantly different in terms of their fits to experimental data as compared to the unreduced full models (p=0.12-0.99, ANOVA chi squared test). Both the pseudovirus mutation and the mutation mismatch models fit training data equally well. The mismatch model, however, gave substantially more accurate predictions of data against each variant pseudovirus when trained on data against all other pseudoviruses (Fig. S4). Accuracy of prediction of data from each serum when models are trained on other sera from the same infecting/vaccine variant was comparable across both the models. Based on these findings, and because the mutation mismatch models had fewer variables, we chose BIC-reduced mismatch mutation models as our best infecting/vaccine variant specific models.

Each vaccine or infecting variant-specific mismatch mutation model fit the experimental data well, but with varying accuracy (coefficient of determination R^2^ = 0.28-0.72) (Fig. 3B). The least accurate predictions were for Delta (R^2^= 0.53) and Lambda (R^2^=0.28) infection sera, but even in those cases, the model fit values were highly significantly correlated with observed titers (Pearson correlation test p < 2.2 × 10^−16^ and 4.8 × 10^−7^, respectively). The low accuracy for Lambda sera was the result of outlier points coming from a single unusual Lambda-infection serum (S19919) that despite having a high autologous ID_50_ = 753, did not neutralize any of the 7 heterologous variants tested (Table S2); if this outlier serum was excluded, a much better R^2^ = 0.55 was obtained. Each infecting/vaccine variant-specific model offered significantly better fits than a model based on autologous titers alone (p = 0.0104, 1.23 × 10^−7^ and < 2.2 × 10^−16^ for sera from Lambda infection, Delta infection, and from each of other infecting/vaccine variants, respectively, using ANOVA Chi-squared test), thus indicating that sequence variables were critical for accurate modeling of the heterologous titer data.

These models allowed identification of sequence variables that significantly impacted serum cross-neutralization resulting from each infecting variant and from mRNA-1273 vaccination (Fig. 3C). Each model had a unique combination of significant sequence mutations with unique coefficients, suggesting that the heterogeneity in cross-neutralization profiles across each vaccine/infecting variant derives from reliance on different mutations with varying magnitudes. While all models had one or more significant RBD mutations, the models for prototype, Beta and Gamma infection sera also showed significant dependence on NTD mutations. The mismatch at the RBD site 371, which only existed for test variants BA.1, BA.1 + A_BA.1_484K and BA.2 when tested against each serum from infecting/vaccine variant in our dataset, made the strongest contribution to cross-neutralizing resistance (0.6-1.0 Log_10_ or 4-10-fold decrease in heterologous ID_50_ titers), consistent with previous reports showing the site-371 mutations cause resistance to all classes of RBD NAbs^10,21^. The next strongest variable was a mismatch between infecting/vaccine and test variants at site 484 that led to 0.6-0.8 Log_10_ drop (4-6-fold) in heterologous titers for mRNA-1273 vaccination sera and Prototype-, Alpha- and Delta-infection sera that all possessed the ancestral E484. In contrast, the models for Beta, Gamma and Iota infecting variants that contained K484 showed no significant dependence on a mismatch at 484, indicating that their immunodominant responses targeted other sites^34^. Interestingly, Lambda-infection sera, despite having an E484 in the infecting variant, did not show resistance at site 484, consistent with the equivalent neutralization of Delta pseudovirus with or without the E484Q mutation (Fig. 2B). This suggests that the Lambda variant may have shifted immunodominance away from the 484 site in the class 2 epitope^42^.

For Alpha, Beta and Gamma-infection sera, our models suggest that the novel epitope created by a mutation at site 501 (N501Y) was immunodominantly targeted, contributing 0.5-1.0 Log10 or 3-10 fold drop in heterologous titers against pseudoviruses lacking N501Y. Comparing the experimental impact of N501Y in the D614G backbone, we found that as predicted, Alpha and Gamma-infection sera showed improved titers against D614G+N501Y as compared to D614G, while no impact was found for mRNA-1273 and Prototype infection sera (Fig. S5, Table S3). While a few Beta-infection sera had elevated titers against D614G+N501Y versus D614G, as predicted, the magnitude of this difference was much lower than predicted and most sera did not show any effect, suggesting that the impact of this mutation may be dependent on the context of the pseudovirus. Amino acid mismatches at sites of the novel mutations in Delta, L452R and T478K, were also associated with significant resistance, suggesting targeting of these novel mutations by neutralizing responses induced by Delta infection. Significant resistance to mutations at site 478, but only minor resistance to mutations at site 452, was found for Delta-infection sera using the deep mutational scanning (DMS) approach^35^. Overall, these models highlight the significant resistance mutations and quantify their relative impact for serum cross-neutralization specific to each infecting/vaccine variant, and provide clues about the potential immunodominant epitopes associated with each infecting/vaccine variant.

These models also allowed delineation of which specific mutations underlie the level of heterologous neutralization resistance of a given variant against sera from people infected by other variants (Fig. 4, S6). For example, for Beta- and Gamma-infection sera, the 2.1-2.4-fold reduction in titers against Alpha pseudovirus is solely attributable to the deletion at site 144 in Alpha, while the 30-fold resistance of Alpha to Delta-infection sera could be traced to the mismatch between Alpha and Delta at sites 452 and 478. Although partial resistance of Kappa pseudovirus to both Alpha- and Gamma-infection sera was attributable to mismatch at site 501 (ancestral N for Kappa, and Y for the infecting variants), the remaining resistance to the Alpha-infection sera came from the E484K mutation in Kappa and to the Gamma-infection sera from the L452R mutation. These results highlight the fine specificity in the targeting of distinct epitopes by NAbs induced by different infecting variants.

**Figure 4:**
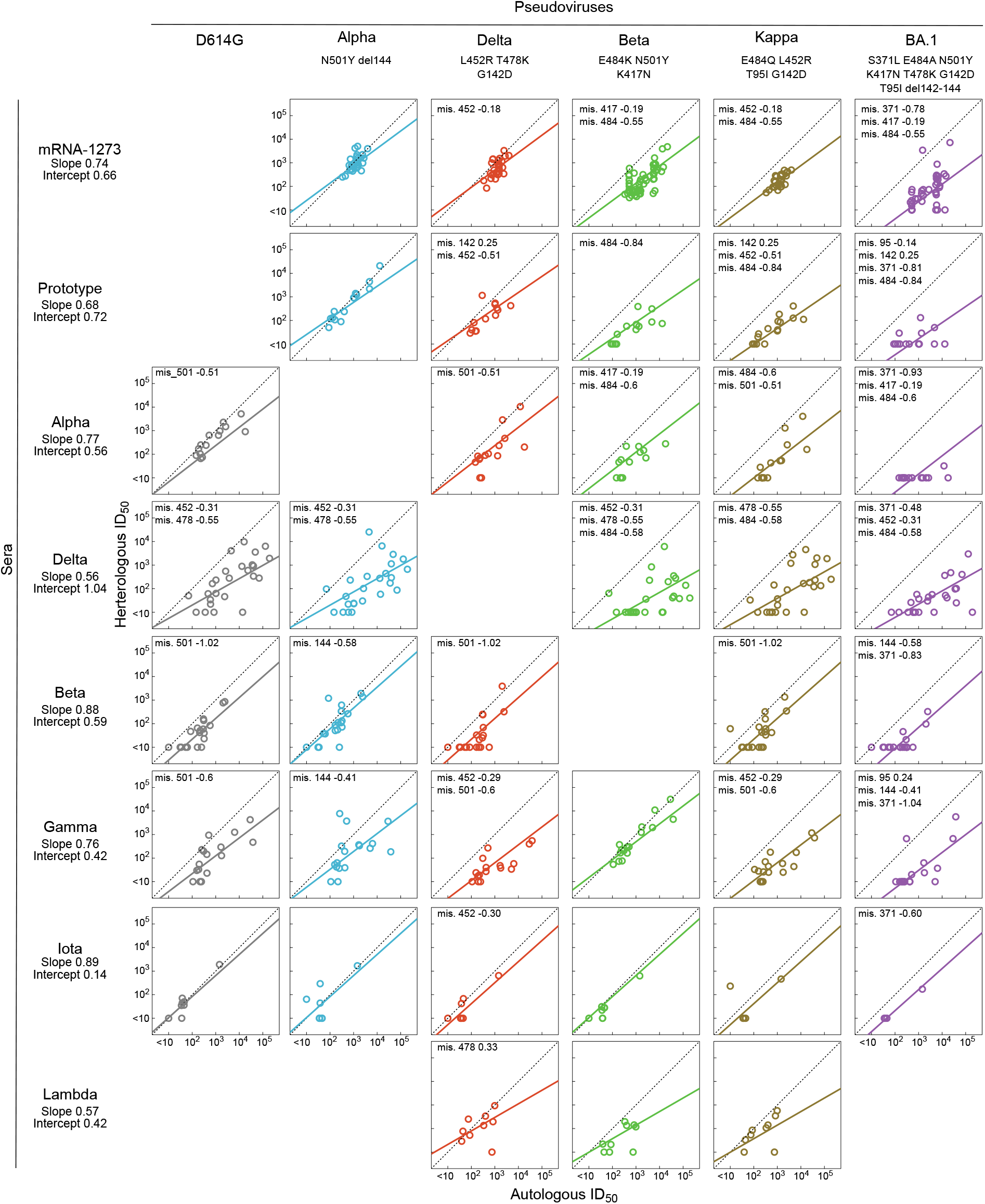
Variant-specific model fits per infecting/vaccine (rows) and test variant (columns). Heterologous ID_50_ titers (vertical) are plotted against autologous titers (horizontal). Dotted line indicates identity and colored solid lines indicate model fits (Fig. 3C). Sequence mismatch variables and their model coefficients (*e.g*. “mis. 501”) for a given pair of infecting/vaccine variant and pseudovirus are shown in top left, and the intercept and coefficient of Log_10_ autologous ID_50_ (“slope”) are shown below each infecting variant.

Curiously, mismatch at three sites, 142, 95 and 478, was associated with significant enhancement of 0.2-0.3 Log10 (1.6-2-fold) for cross-neutralizing titers against D614G, Gamma and Lambda infection sera (Fig. 3C). Such cross-neutralization enhancement associations with heterologous mutations could arise from improving biophysical interactions between NAbs and Spike (*e.g.* improving exposure of the targeted epitope), or from reducing binding to competing non-neutralizing antibodies that hinder binding of neutralizing antibodies^16^. Alternatively, positive coefficients for mutational variables could arise in our modeling framework to compensate for the effects of other variables for certain variants. Whether this phenomenon is a modeling artifact or has a biological underpinning remains to be tested.

### A unified statistical model captures heterogeneous cross-neutralization profiles across infecting and vaccine variants

Given the patterns of mutational dependence that emerge across the infecting or vaccine variant-specific models, we next investigated whether a unified pan-variant model could account for these patterns. Such a unified model could potentially capture the overall relationships between the sequence of a variant and the cross-neutralizing responses it induces, and could predict responses resulting from infection or vaccination by novel variants.

Since it was not clear *a priori* what the functional form for such a unified model should be, we explored four models that included different types of sequence variables, and interactions thereof, in addition to the autologous ID_50_ titers. The simplest included only amino acid mismatch variables between infecting/vaccine and test variants at specified sites, similar to above infecting/vaccine variant-specific models. The second included a linear combination of the mismatch variables and sequence variables that captured the mutations of the infecting/vaccine variant, based on the rationale that such a model would better capture the differential dependence on mutations in test pseudoviruses based on the infecting variant in question. The third used same variables as the second, but also included interactions between mismatch and infecting variant mutation variables at the same site, allowing added flexibility in estimating the impact of mismatch variables conditional on the mutations in the infecting/vaccine variant at the same site. The fourth extended these interactions to be present between mismatch variables and infecting variant mutational variables at *any* two sites, with the rationale that if the infecting variant has a particular mutation at a given site, it can potentially drive the responses to target another site. The complexity increases across these models, with the first model having a maximum of 9 sequence variables, the second 20, the third 31 and the fourth 119. Thus, to identify robust minimal models, we used model selection based on AIC or BIC as described in STAR Methods. Equivalent models using the mutational variables for the test variant instead of the infecting variant were also evaluated, but these did not show improved performance.

The different models and model selection strategies were evaluated for their accuracy of prediction using cross-validation to explore the potential of such models to predict cross neutralization profiles for novel future variants (Table S4). For this, each variant’s data either as infecting variant or as pseudovirus was removed from the training data, and the models fitted to such training data were evaluated for accuracy of prediction on the removed data stratified by either having the variant in question as infecting/vaccine variant or as pseudovirus; this process was iterated over all variants in the dataset. The third model (with forward BIC model selection) performed the best for both infecting variant and test variant cross-validation (Fig. 5A, Table S4). The performance of this statistical model was also compared to non-linear machine learning regression algorithms using the same variables (Extreme randomized forest with and without feature selection using minimal Redundancy Maximum Relevance (mRMR) and nested cross-validation (STAR Methods)). While the machine learning models provided slightly better cross-validation prediction accuracy (the average coefficient of determination R^2^ = 0.61 versus 0.58), the third statistical model had the notable advantages of simplicity and interpretability, with coefficients quantifying the relative impact of each input variable. Thus, we retained this third model as our best unified model.

**Figure 5:**
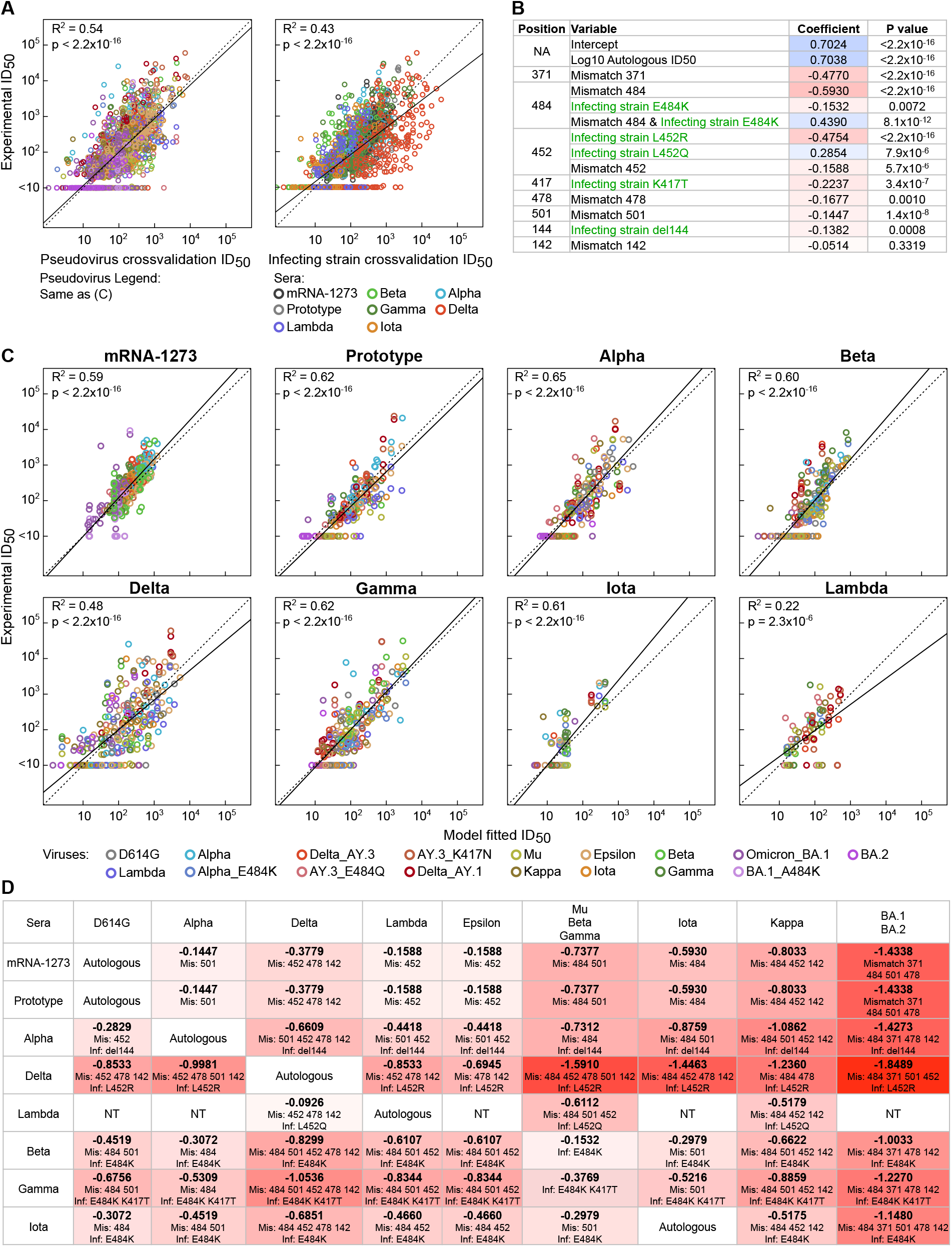
Unified model for cross-neutralization of pan-infecting/vaccine variant sera. (A) Experimental ID_50_ titers (vertical) are plotted against predicted ID_50_ titers (horizontal) for pseudovirus cross-validation (left) and infecting/vaccine variant cross-validation (right). Left subpanel point colors indicate pseudoviruses, per legend in (C), and right subpanel point colors indicate infecting/vaccine variants. Dotted line is identity and solid line is linear fit. (B) Coefficients of the best-fit unified model with their statistical significance calculated using Student’s t-test. Same formatting as Fig. 3C, except infecting variant mutation variables are shown in green. (C) Experimental ID_50_ titers (vertical) are plotted against best-fit model ID_50_ titers (horizontal) with same formatting as Fig. 3B. (D) Cells indicate the net contribution of sequence variables in the unified model to heterologous Log_10_ ID_50_ for each pseudovirus (columns) by sera from each infecting/vaccine variant (rows) with contributing sequence variables noted below. Model predictions for Mu, Beta, and Gamma pseudoviruses, and for BA.1 and BA.2 pseudoviruses, were identical across all infecting/vaccine variants in our dataset and are thus combined.

Fitting the third model on the full training dataset revealed how it could capture the heterogeneous cross-neutralization profiles across all infecting and vaccine variants (Fig. 5B-C, Fig. S7-S8). The mismatch variables at sites 371, 484, 452 and 501, which were most frequently associated with resistance across several infecting/vaccine variant-specific models, were also found to be significantly resistant variables in this unified model (Fig. 5B). In addition, mismatch at site 478, associated with strong resistance to Delta infection sera (Fig. 3C, 4), also contributed significant resistance to the unified model, while mismatch at site 142 contributed insignificant resistance (−0.05 Log_10_, p = 0.33). Infecting variants having the non-ancestral mutations L452R (Delta), E484K (Beta, Gamma, Iota), K417T (Gamma) and deletion at site 144 (Alpha) had significant penalties and those with L452Q (Lambda) had a significant positive contribution. These terms operate regardless of the test variant and thus capture the overall infecting variant-specific trends such as the strong reduction of heterologous titers for Delta infection sera as compared to autologous titers, and the relatively lower reduction of heterologous titers for Lambda-infection sera (Fig. 2A). A significant interaction was identified at site 484, indicating that the impact of mismatch in infecting and test variant mutations at site 484 was dependent on the mutation at site 484 in the infecting/vaccine variant. For E484 infecting variants, the net effect of mismatch at site 484 was 0.59 Log_10_ resistance, while for K484 infecting variants, the effect was 0.15 Log_10_ resistance, consistent with the patterns of site 484 resistance seen above for infecting/vaccine variant specific models (Fig. 3C). A notable departure from infecting variant-specific models was the absence of resistance of mismatch at site 144 for Beta and Gamma-infection sera (Fig. 3C, 4, 5B). This mismatch is present when these sera were tested against Alpha, Mu, and BA.1 pseudoviruses (Fig. 4, S6) and when compared with corresponding fits for the unified model (Fig. 5C, S7, S8), the contribution of site 144 mismatch is replaced by that from site 484 mismatch (Alpha, BA.1) and the contribution for infecting variant with E484K (Mu). This suggests that at least from a modeling standpoint, the roles of sites 144 and site 484 variables were effectively interchangeable, and in the context of a minimal model constructed using stringent model-selection criteria, the site 484 variables may suppress the site 144 variables.

The unified model provides a useful mathematical representation of the cross-neutralization profiles across all infecting/vaccine and test variants since infecting/vaccine variant-specific model fits were only marginally better (Fig. 3B, 5C). Because infecting variant mutational variables are included in the third unified model, the heterologous titers are not symmetric upon exchange of infecting and test variants as would have been the case if only mismatch terms had been included. With the exception of two clusters of related test variants (Mu, Beta and Gamma; and BA.1 and BA.2) (Fig. 5D), this single unified model produces highly heterogeneous sequence penalties for each combination of infecting/vaccine variant and test variant, thus capturing the observed cross-neutralization heterogeneity.

### Accurate prediction of novel VOC and vaccine cross-neutralization

The accuracy of the unified model in fitting the cross-neutralization data suggests that it can capture the basis of infecting/vaccine variant-specific patterns of cross-neutralization of SARS-CoV-2 variants. To test this, we examined the ability of this model to predict cross-neutralization titers from novel vaccine and infecting variants as well as against novel test variants (Fig. 6, Table S5). First, we tested prediction accuracy for sera from a primary 2-dose novel vaccine based on the Beta variant, mRNA1273-351, and found significant agreement between model predictions and experimental titers (Fig. 6A, R^2^=0.43, p = 6.4×10^−16^ Kendall tau rank test). There was a systematic trend in our predictions being lower than experimental titers, but this difference was small (~0.2 Log_10_ or 1.6-fold). When the Beta-infection sera model was used for prediction, a lower but still significant accuracy was found (R^2^= 0.20, p = 3.1×10^−5^ Kendall tau rank test). Thus, while Beta mRNA vaccination led to similar responses as primary Beta-infection convalescent sera, the unified model was better than the Beta infection-specific model at predicting these responses. Since the Beta-infection model was trained only on Beta-infection sera data, while the unified model was trained on the full dataset, the reduced accuracy for the Beta-infection specific model in predicting Beta-mRNA vaccination data could arise due to over-fitting of patterns in Beta-infection sera data. We also predicted heterologous neutralizing titers for three primary Omicron BA.1 infection sera and found them to be significantly accurate (Fig. 6B, R^2^ = 0.59, p = 3.4×10^−6^); an underestimation of experimental titers by ~0.2 Log_10_ was again found. These examples demonstrate that the unified model can accurately predict cross-neutralizing activity induced by novel variants in the setting of primary vaccination or infection.

**Figure 6:**
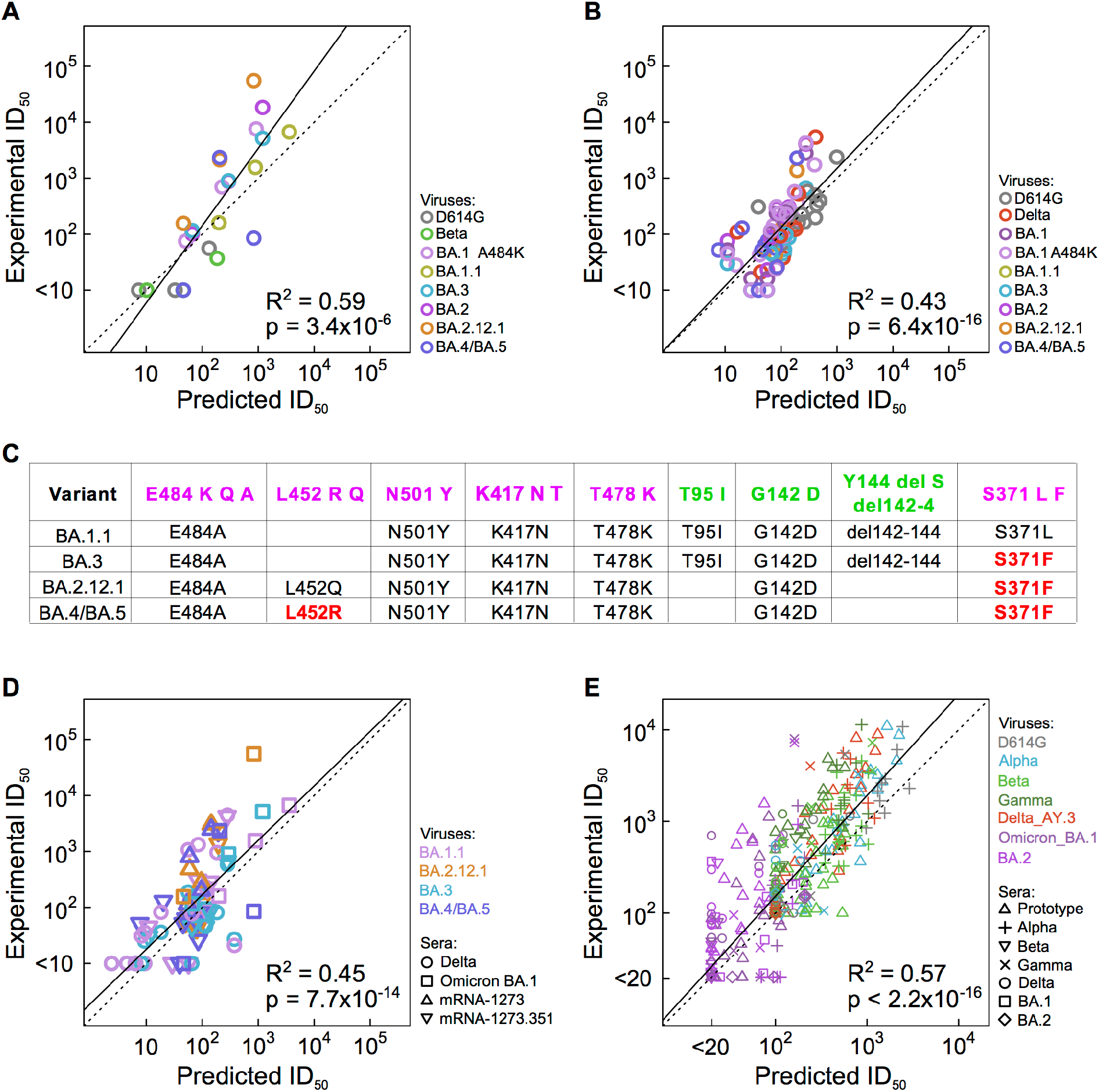
Accurate prediction of holdout cross-neutralization data. Model predictions (horizontal) and experimental ID_50_ titers (vertical) are shown for Omicron BA.1 primary infection sera (A), Beta mRNA (mRNA-1273.351) vaccine sera (B), and primary infection sera from an independent dataset^37^ (E). Panel (C) shows sequence of novel pseudoviruses BA.1.1, BA.3, BA.2.12.1 and BA.4/BA.5 at the sites used by the model, and panel (D) shows prediction of cross-neutralization titers of these novel pseudoviruses. For panels (A), (B), (D) and (E), point colors indicate pseudoviruses, and for panels (D) and (E), different symbols indicate sera from different infecting/vaccine variants. Dotted lines indicate identity and solid black lines indicate linear fits. Coefficient of determination (R^2^) and two-sided p-values from Kendall tau rank test are shown.

We next investigated the prediction ability of our models in the context of four holdout pseudoviruses – BA.1.1, BA.3, BA.2.12.1 and BA.4/BA.5 (baseline BA.4 and BA.5 have the same Spike sequence). BA.1.1 differs from BA.1 at a single Spike site (R346K), while BA.3 and BA.1 have differences at 7 sites (214, 371, 405, 496, 547, 856, 981) (Fig. S2). BA.2.12.1 differs from BA.2 at two sites (L452Q and S704L) and BA.4/BA.5 differs from BA.2 at 5 sites (deletion at 69-70, L452R, F486V and R493Q). At the sites used in our model, BA.1.1 was identical to BA.1, and BA.3 was different at a single site (F371), while BA.2.12.1 and BA.4/BA.5 differ from BA.2 at site 452 (Fig. 6C). These four viruses were tested against mRNA1273-351 vaccinated and primary BA.1 infection sera from above, in addition to four mRNA1273 vaccinated and 15 Delta infection sera from our training dataset. Our model predicted these neutralizing titers with significant accuracy (R^2^ = 0.43, p = 7.7×10^−14^), with again a slight underestimation of titers of ~0.2 Log10.

We also tested the prediction accuracy of our model on an independent dataset published by Van Der Straten et al.^37^, including ID_50_ neutralization titers of 64 sera from primary infections with one of 7 VOCs (D614G, Alpha, Beta, Gamma, Delta, BA.1 and BA.2) tested against 8 VOC pseudoviruses (all the above, except Beta, which was tested as two variants, L242H+R246I and Δ242-246)^37^. Our model predicted these titers with significant accuracy (R^2^ = 0.57, p < 2.2×10^−16^; Fig. 6D). It also predicted the low breadth of BA.2 infection sera, correctly predicting all 24 of 28 experimental heterologous ID_50_ titers that were below threshold, and 2 of 4 positive titers (accuracy = 0.93, odds ratio = ∞, p = 0.0159, Fisher’s exact test).

### Third mRNA-1273 dose lowers the impact of resistant mutations

We next investigated whether our model, trained on primary infection or primary 2-dose vaccination series, could predict heterologous neutralization for 3-dose mRNA-1273 boosted sera. We tested 26 boosted sera from the clinical studies DMID 21-0002 and DMID 21-0012^44^ for neutralization against D614G and 7 variant pseudoviruses (Beta, Delta, BA.1, BA.1.1, BA.3, BA.2.12.1 and BA.4/BA.5) and one mutant pseudovirus (BA.1 + A_BA.1_484K) (Table S6). Neither the 2-dose mRNA-1273 specific model nor the unified model accurately predicted the heterologous titers for 3-dose sera (R^2^ < 0.0). To understand this, we first compared cross-neutralizing titers of 2-dose and 3-dose sera as a function of autologous titers. While D614G and Delta pseudovirus titers showed no difference between the 2- and 3-dose sera, the 3-dose mRNA-1273 sera exhibited disproportionately better cross-neutralization of Beta, BA.1, BA.2.12.1 and BA.4/BA.5 pseudoviruses than 2-dose sera, even after correcting for elevated autologous titers (Fig. 7A). Thus, boosting with mRNA-1273 led to improved cross-neutralization, especially of Omicron lineage variants, which is consistent with previously observed better recall and/or maturation of cross-reactive B cells, and induction of novel B cell clonal lineages with higher targeting of more conserved epitopes post the third dose^45–47^.

**Figure 7:**
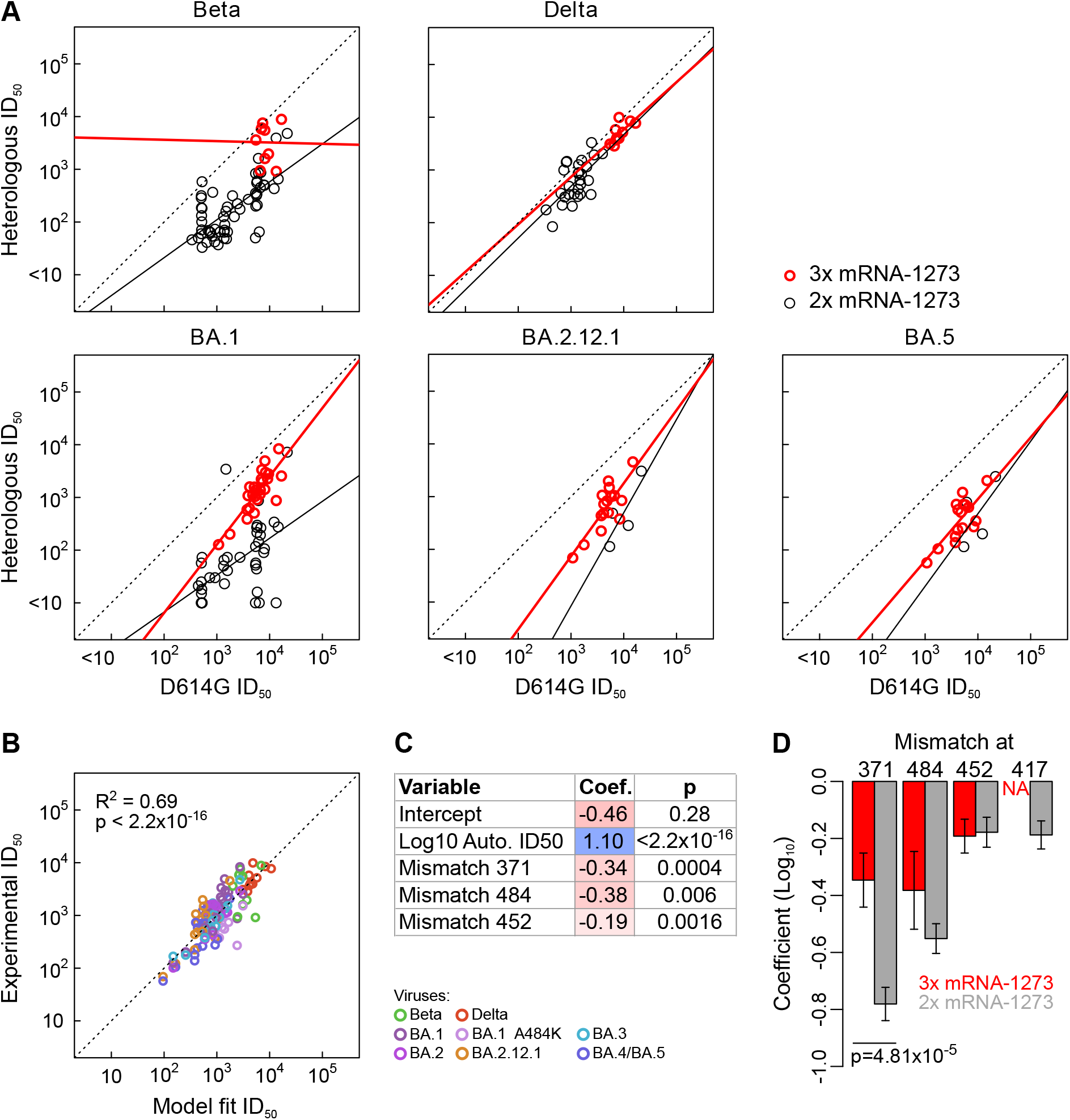
Comparison of 3-dose to 2-dose mRNA-1273 vaccinated serum cross-neutralization. (A) Heterologous ID_50_ titers for indicated pseudoviruses (vertical) are plotted with D614G titers (horizontal) for sera post 3-dose (red circles) and post 2-dose (black circles) mRNA-1273 vaccination. Dotted line is identity, and black and red solid lines show linear fits for 2-dose and 3-dose Log_10_ titers, respectively. (B) Plot of experimental (vertical) versus model fitted (horizontal) heterologous ID_50_ titers for 3-dose sera, with point colors indicating pseudoviruses. (C) Variable coefficients for the 3-dose mRNA-1273 model and their significance using Student’s t-test. Color scale as in Fig. 3C. (D) Comparison of coefficients for sequence variables between 3-dose (red) and 2-dose (grey) models. Error bars indicate twice the standard error (S.E.). Statistical significance for difference in coefficients was calculated using Z-test, and only the mismatch at site 371 had p < 0.05.

To further explore the broader neutralization of post 3^rd^ mRNA-1273 dose sera, we modeled heterologous titer data using the same strategy employed for post 2^nd^ dose sera, and found similarly significantly accurate fits to the experimental data (R^2^ = 0.69, p < 2.2×10^−16^ Kendall tau rank test; Fig. 7B). In this model, mutations away from ancestral Wuhan-1 at only 3 positions contributed to resistance: 371, 484 and 452 (Fig. 7C). These mutations also contributed to significant resistance for the 2-dose mRNA-1273 model, but the magnitude of resistance for the 2-dose model was significantly higher for site 371 (difference = 0.43 Log_10_ or 2.7-fold; p = 4.8 × 10^−5^, one-sided Z-test) and higher, but not significantly, for site 484 (difference = 0.17 Log_10_ or 1.48-fold, p=0.12). Predicted resistance due to mutations at site 452 in the Delta, BA.2.12.1 and BA.4/BA.5 pseudoviruses was comparable between the two models. While the 2-dose model also showed significant resistance from mutations at site 417 (Fig. 3C), the available data for 3-dose sera precluded independent assessment of this mutation as it was always accompanied by a mutation at site 484 for the pseudoviruses tested. Overall, these analyses indicate that the higher neutralization breadth of the 3-dose mRNA-1273 vaccinated sera as compared to the 2-dose sera derives from improved recognition of resistance mutations at sites 371 and 484.

## DISCUSSION

We demonstrated that the cross-neutralization titers of sera from primary infection with any variant or from primary 2-dose mRNA-1273 vaccination could be accurately modeled using autologous titers and sequence variables of infecting/vaccine and pseudovirus variants. The heterogeneity of cross-neutralization profiles seen across different infecting or vaccine variants and across pseudovirus variants arises due to the commonality or difference of mutations at specific sites. We could resolve the impact of specific mutations in part due to the high similarity of cross-neutralizing responses across hosts infected or vaccinated with the same variant, suggesting that each SARS-CoV-2 variant reproducibly induces specific immunodominant cross-neutralizing responses. However, such responses vary across infecting/vaccine variants, highlighting that the variants differ not only in their neutralization sensitivity/resistance to monoclonal NAbs and convalescent or vaccinated sera^5,10,21,40^, but also in the induction of NAb responses post infection or vaccination.

Our modeling identified distinct mutations associated with this variability and quantified the magnitude of their impact. Previous studies have analyzed this using DMS^16,34,35^ and antigenic cartography^36,37^, each providing important perspectives. DMS-based approaches identify single mutations that impact serum or antibody neutralization and ACE2 binding, including those not sampled by current or previous variants, but they are limited to analyzing a few background Spike constructs and cannot identify the combined effect of multiple resistance mutations. While antigenic cartography provides an informative visualization of the relationship of neutralization profiles of sera from people infected or vaccinated with particular variants among themselves and relative to the pseudovirus variants, it cannot readily identify the specific mutations that underlie these relationships nor quantify their relative impacts. By identifying significant mutations impacting cross-neutralization and quantifying their relative impacts in the context of circulating variants, our complementary modeling strategy provides additional insights into the antigenic and immunogenic variability of SARS-CoV-2 variants.

Differences in serum cross-neutralization profiles induced by different variants imply that there are variant-specific differences in the nature and presentation of immunodominant neutralizing epitopes, as shown by DMS studies of Omicron variants^48^. Since such epitope properties are ultimately governed by the sequence of the variant, this implies the existence of a well-defined relationship between the sequence of the infecting/vaccine variant and the induced cross-neutralizing profiles. Indeed, the accurate modeling and prediction of cross-neutralizing data by our unified model supports the existence of this relationship. However, it is remarkable that this relationship can be accurately quantified using only sequence variables at 8 sites along with autologous ID_50_ titers (Fig. 5B). The biological underpinnings of this could potentially arise from several observations. First, only a few distinct neutralizing antibody classes are immunodominant in the population level response to SARS-CoV-2 variants^16,40–42^, suggesting that only a limited number of epitopes on Spike are relevant for most of the serum cross-neutralization in the population. Second, distinct variants may, through the nature of the epitopes they present, modulate the relative propensity of induction of each of these few immunodominant NAb classes. For example, SARS-CoV-2 variants like Beta^34^ and Omicron that have evolved to escape preexisting neutralizing immunity could potentially induce alternative immunodominant NAb responses due to their mutations abrogating the epitopes of previously common NAb responses in the population. Third, certain mutations can impact multiple immunodominant NAb classes thereby reducing the number of Spike sites that potentially modulate cross-neutralization profiles. Experimentally validated examples of such are sites 371 and 452 used in our models^10,21^, and sites 346, 460 and 486^7,9,10,16^ that are often mutated in emerging Omicron subvariants.

While our study used historic data, it provides proof-of-concept that our approach can be applied for accurate modeling of more contemporary data on Omicron and recombinant lineages and from an increasingly more hybrid immune population. If successful, such approaches could help with anticipating emergence of novel neutralization resistant variants and designing of next-generation SARS-CoV-2 vaccines.

## LIMITATIONS OF THE STUDY

Our models are based on neutralizing responses induced by primary infection with known specific variants and by primary 2-dose mRNA vaccination. However, such scenarios are rare in the current stage of the pandemic, with most of the global population having complex histories of multiple vaccinations and one or more infection(s). The observed difference in neutralizing activity between 2 and 3 doses of mRNA-1273 (Fig. 7) suggests that multiple antigenic exposures can lead to strikingly different neutralizing responses, as shown for multiple exposures to Omicron^48^. While this is an important limitation of this study, our sequence-based modeling of neutralizing responses can be adapted to analyze current and future scenarios, to better understand: vaccine responses based on new variants, the impact of circulating mutations on vaccine efficacy to inform future vaccine design efforts, and the role of emergent population level neutralizing immunity on the continuing evolution of SARS-CoV-2 and the epidemiology of the pandemic. Furthermore, our current models of primary infection or primary vaccination could prove useful in understanding how hybrid immunity from multi-variant vaccines, vaccine breakthrough or multiple infections can be modeled as a function of responses expected from each exposure, and thus facilitate insights in the complex immunology of multi-antigen exposures (*e.g*., original antigenic sin, primary addiction, back-boosting, etc.).

Another limitation is that our model is based on recurrent mutations that arose in multiple previous variants. While some of these sites continue to vary between major Omicron sublineages, such as acquisition of L452R (BA.5 sublineages), L452M (BA.2.3.20), E484R (BA.2.3.20) and Y144- (XBB.1), other mutations chosen based on older variants for our models may not be particularly relevant for the ongoing diversification of Omicron sublineages. This is consistent with recent work showing that the backbone of Omicron variants has allowed the exploration of different antibody escape mutations that were deleterious in earlier variants since they reduced ACE2 binding, but are not in the Omicron backbone due to epistatic effects^13,39,49^. As the landscape of immunologically relevant mutations shifts, our current models will have to be updated to better capture such mutations. Nonetheless, our modeling strategy provides a proof of concept regarding the value of sequence-based modeling, and a template for how newer models can be developed.

## Acknowledgements

Figure S1 was based on global sequences shared through GISAID since the start of 2021, and we thank all data contributors for obtaining the samples and sequences to enable a global view of viral evolution. We thank Haili Tang and Wenhong Feng for producing and generating the plasmids and viruses; and Charlene McDanal, Jin Tong, and Lily Daniell for assay support. We greatly appreciate John Hural, Agatha Jassem, Yoshihiro Kawaoka, Vic Veguilla, Florian Krammer, Viviana Simon, Richard Webby, Victor Corman, Christian Drosten and Paul Chris Roberts and the DMID 21-0002 and DMID 21-0012 study teams for providing valuable vaccine and infection sera for the study. We thank Christine Posavad for assistance in Infectious Disease Clinical Research Consortium (IDCRC) manuscript approval process. This work is supported by the Los Alamos National Laboratory lab-directed R&D exploratory research project (LANL LDRD 20220399ER) (K.W., B.K.), NIAID-NIH SARS-CoV-2 Assessment of Viral Evolution (SAVE) program (#75N93019C00050) (X.S., D.M., B.K.) and NIAID-NIH Design and development of Pan-betacoronavirus vaccine (P01 AI158571) (K.W., B.K.). This study was also supported by the Infectious Diseases Clinical Research Consortium through the National Institute of Allergy and Infectious Diseases, part of the National Institutes of Health, under award number UM1AI148684. The content is solely the responsibility of the authors and does not necessarily represent the official views of the National Institutes of Health. Pseudovirus neutralization (PsVN) assay data provided by Duke was supported in part with support from the NIAID Collaborative Influenza Vaccine Innovation Centers (CIVICs) contract 75N93019C00050.

## Author contributions

Conceptualization: K.W., B.K.; Methodology: K.W., X.S., J.T., D.C.M, B.K.; Software: K.W.; Validation: K.W., X.S., J.T., D.C.M., B.K.; Investigation: K.W., X.S., B.K.; Formal analysis: K.W., J.T., B.K.; Resources: X.S., B.G., J-C.M., D.C.M.; Writing - Original Draft: K.W., X.S., J.T., D.C.M., B.K.; Writing - Review & Editing: K.W., X.S., J.T., B.G., D.C.M., B.K.; Supervision: J.T., D.C.M., B.K.; Funding Acquisition: K.W., D.C.M., B.K.

## STAR Methods

### Clinical samples

Serum samples from individuals that received the Moderna mRNA-1273 or mRNA-1273.351 vaccine or were infected with the prototype D614G or other SARS-CoV-2 variants are used in this study. All samples, including those from vaccine recipients were collected in clinical studies with information listed in Table S1. Convalescent sera are assumed but not confirmed first infections. Institutional review board (IRB) approvals were obtained for each study site. All participants were given and signed the informed consent form. The use of the specimens for the research conducted in this manuscript is covered under IRB protocols Pro00093087 and Pro00105358. Three-dose mRNA-1273 samples and data were provided by DMID 21-0002 and DMID 21-0012 study teams and these studies were supported by Advarra IRB. DMID 21-0012 samples are from Lyke et al.^44^ and DMID 21-0002 samples were provided by the DMID 21-0002 study team in support of the partial pseudovirus neutralization (PsVN) assay validation for the Beta variant of SARS-CoV-2.

### Neutralization assays

A lentiviral vector pseudovirus neutralization assay was used to quantify neutralizing activities of the serum samples. Pseudoviruses were prepared, titrated and used for measurements of neutralizing antibodies essentially as described previously^5^. Briefly, Spike mutations were introduced into the VRC7480 plasmid (an expression plasmid encoding codon-optimized full-length spike of the Wuhan-1 ancestral sequence that was provided by Drs. Barney Graham and Kizzmekia Corbett at the Vaccine Research Center, National Institutes of Health (USA) by site-directed mutagenesis. Pseudoviruses were produced in HEK293T/17 cells by co-transfection of a spike plasmid, a lentiviral backbone plasmid, and firefly Luc reporter gene plasmid, and a TMPRSS2 plasmid. Pseudovirus culture was collected after 2 days of incubation and titrated as described previously ^5^.

Neutralization assays were performed with pre-titrated pseudoviruses using a 293T cell line that has been engineered to stably express ACE2 (293T/ACE2-MF) as previously described ^19^. Spike-pseudovirus was incubated with serially diluted serum samples in duplicate for 1 hr before cells were added to the mixture. After 71-73 hrs of incubation, cells were lysed using Bright-Glo luciferase reagent and luminescence was measured using a GloMax Navigator luminometer (Promega). Neutralization titers are the inhibitory dilution (ID) of serum at which relative luminescence units were reduced by 50% (ID_50_) compared to virus control wells.

#### Down selection of Spike sites for modeling

Sequences used for pseudovirus variants are shown in Fig. S2. While the infecting variant information for the serum samples was available, the exact sequence of infecting virus was not. Thus, we assumed that the infecting variant matched the corresponding pseudovirus variant. To ensure statistical robustness of our models, we restricted our selection to all sites that were mutated away from the prototype in at least 3 wildtype variants in our dataset (counting the closely related Delta AY.1, AY.2 and AY.3 as a single variant) (Fig. S2B). This resulted in 10 sites after excluding D614G, which was present in all pseudoviruses. From these 10 sites, we identified a set of 8 sites – 5 in RBD (417, 452, 478, 484, 501) and 3 in NTD (95, 142, 144) – that had mutations that when expressed as binary variables (1 if mutated, 0 if prototype) were linearly independent. We note that this choice of sites is not unique, and we were guided by trying to select sites in commonly targeted neutralizing antibody epitopes in RBD and NTD that had common antibody resistance mutations^40–42^. With the origin of BA.1 and BA.2, it has been shown that the dramatic resistance to RBD neutralizing antibodies of these variants can be tracked predominantly to S371L and S371F, respectively^10,21^. Based on this and its linear independence to the above sites, we added 371 to the list of chosen sites. These 9 chosen sites had a total of 16 mutations in our dataset.

Due to redundancy between chosen sites and other sites, our models cannot distinguish between the effect of a given downselected mutation and that of remaining mutations that are linearly dependent on it. For example, the deletion at sites 69 and 70, which we left out, co-occurs with a deletion at 144 in Alpha and BA.1, and thus, the impact of deletion at 144 found in our models could be a surrogate for neutralization resistance coming from the deletion at 69-70. Five mutations (K417T, L452Q, Y144S, S371L, S371F) were found in only one variant, and thus, any importance attributed to these mutations could come from any other variant-specific mutations that we did not include in our sequence variables. We also removed data for pseudovirus Delta AY.2 as it showed highly correlated titers with AY.3+K417N (Fig. S3), and had exactly identical mutations to AY.3+K417N at the above 9 sites (Fig. S2). While titers against Delta AY.1 were also highly correlated with AY.3+K417N (Fig. S3), it had a T95I mutation that was not shared with the latter, and thus, we retained Delta AY.1 pseudovirus data in our final set.

#### Statistical and machine learning modeling

All statistical models were developed using R (version 3.3.3 GUI 1.69 for Mac OS)^50^ using the generalized linear model ‘glm’ function. The exact formulae for infecting/vaccine variant-specific models and the four model families for the unified model are shown in supplemental information, together with the summaries detailing the coefficients and significance for each of the models in Figures 3, 5 and 7.

For AIC or BIC based model selection, the ‘step’ function in R was used. For forward selection, the starting baseline model for Log10 heterologous ID50 titers was a linear model of Log10 autologous ID50 titers and an intercept, and giving the full formula as the ‘scope’ parameter in the ‘step’ function. We used ANOVA as implemented in R to calculate the significance of reduction in model fits for AIC or BIC reduced models as compared to the full models for each infecting/vaccine variant. For the smallest two unified models, those with mismatch terms and with mismatch and infecting variant mutation terms, only backward selection, i.e., sequential removal of variables that do not contribute significantly to fitting the training data, was used. For the intermediate sized third model, both forward and backward model selection was used. For the largest model (model #4), only forward selection, i.e., sequential addition of variables that significantly improve fits to the training data was used. All other statistical tests were performed using packages in R. While multiple tests correction preclude this, we have continued to use uncorrected p < 0.05 as the threshold for statistical significance to ensure a more inclusive threshold for identification of important sites.

For machine learning, we used the ExtraTreesRegressor function from the Scikit Learn package^51^ in Python with and without feature prefiltering using the mRMR algorithm^52^ based on our previous success of modeling antibody neutralization titers using this approach^53^. Optimal parameters of the regressor such as the number of estimators (range 20-500) and minimum samples split (range 2-20) and the optimal fraction of top mRMR selected features to be retained for modeling (range 0-100%) were identified using a nested cross-validation approach described below.

#### Cross-validation analyses

For unified statistical models and machine learning models, cross-validation (CV) was used to identify the best models. For each variant the full dataset was separated into a training set and two test sets, and this procedure was iterated over all variants. The first test set for a given variant included all the neutralization data for sera that had this variant as an infecting/vaccine variant. The second test set included all the data points that had this variant or its mutants as the pseudovirus (*e.g*. Alpha and Alpha+E484K). The division in two test sets was done to allow measurement of prediction accuracy of a given model in predicting data for novel infecting/vaccine variants and for novel pseudoviruses that could *a priori* be different. The training dataset for a given variant was all the data that did not fall in the two test sets for the variant. For statistical models, the models were first trained on the training dataset and then applied to either test set to obtain predictions. For machine learning models, since cross-validation is also used to optimize algorithm parameters, this parameter selection step can introduce biased (i.e., artificially more accurate) predictions using conventional CV^54^, which could also hold true for the above CV. Thus, we followed the procedure of outlined by Varma et al.^54^ of using an “internal loop” of CV on the training set for each “external” CV loop to identify optimal parameters. To achieve this in our context, the training dataset for each variant was subjected to the same CV procedure, i.e., dividing the data into internal training and internal test datasets based on the variants available in the training set, with the exception that the two types of internal test datasets were combined. Optimal ML parameters were identified as those that gave the best prediction accuracy on this internal CV, and then these parameters were used to construct models that were evaluated for their prediction of the external test datasets. For both statistical and ML modeling, the predictions for test sets for either the first type (infecting variant CV) or the second type (pseudovirus CV) were compared to experimental titers across all variants using the coefficient of determination R^2^ values. Averaged R^2^ across these two flavors of CV was used as the single metric to rank the different modeling strategies.

#### Comparison to van der Straten et al. data

The heterologous serum neutralizing titers reported in van der Straten et al. were in IU/ml^37^. To facilitate comparison with ID_50_ titers in our study, we converted the IU/ml to ID_50_ titers using the conversion ID_50_ = 10 × IU/ml, as reported by the authors. The sequences of pseudoviruses used in that study also differed slightly from those in our study, *e.g*. Beta, and thus, we calculated sequence variables for model predictions using their reported sequences.

