## Supplementary Information for "Mutational basis of serum cross-neutralization profiles elicited by infection or vaccination with SARS-CoV-2 variants"

**Supplementary Table & Figure Legends:**

**Table S1: Clinical characteristics of analyzed samples (related to Fig. 1-5).**

**Table S2: Neutralization dataset used for model training (related to Fig. 1-5).** While only ID50 titers were used in our models, ID80 titers are also provided. Infecting strain or vaccine information is provided along with patient ID (PTID). Rows are sera and columns are pseudoviruses, with ID50 titers followed by ID80 titers.

**Table S3: ID50 titers against D614G and D614G+N501Y for select sera (related to Fig. 3).**

**Table S4: Cross-validation (CV) prediction accuracy across different modeling strategies for unified models (related to Fig. 5).** Each row indicates a given modeling strategy in the first column, and the second and third columns indicate coefficient of determination ($R^2$) between predicted and observed ID50 titers using either pseudovirus or infecting/vaccine strain CV as described in STAR methods. The last column indicates the averaged $R^2$ between the two CV. Similar models from the same model "family" are colored similarly. "ExtraTrees" is the extremely randomized machine learning (ML) algorithm, and MRMR indicates feature selection using minimal redundancy maximal relevance. All other models are generalized linear statistical models. "Forward AIC" or "Forward BIC" indicates forward model building (i.e. sequentially adding variables) using either AIC or BIC, respectively and all models with "AIC" or "BIC" indicate backward model selection (i.e. removing variables) using AIC or BIC. Models without AIC or BIC in their names are full models using all variables from the corresponding formulae (Text S1), with the exception of the two ML algorithms where AIC/BIC model selection cannot be done. For comparison, we have also included a model using only autologous ID50 titers, and the infecting/vaccine strain specific models similar to Fig. 3. For the latter, the model corresponding to each infecting/vaccine strain was used for prediction of test dataset, and because of this constraint, only pseudovirus CV can be performed.

**Table S5: Holdout dataset used for model validation (related to Fig. 6).**

**Table S6: ID50 titers for sera from 3-dose mRNA-1273 vaccinated participants (related to Fig. 7).**

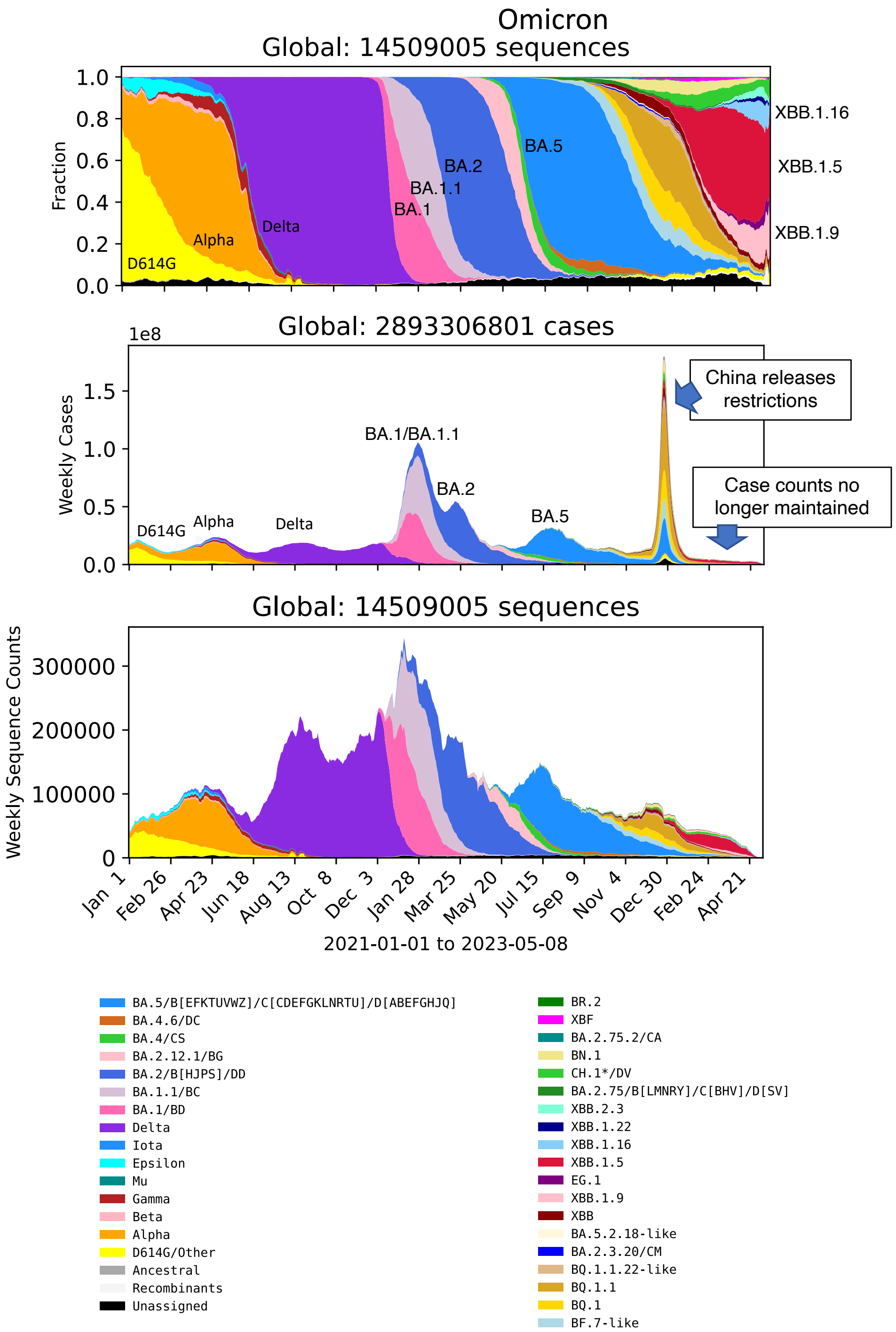


**Figure S1: Major variant transitions have been historically associated with global case waves (related to Fig. 1, 2).** Frequency of major variants forms (top) and predicted weekly number of cases attributable to each variant (middle), and weekly number of reported sequences (bottom), are shown from 1st January 2021 to 12th April 2023. Data for each variant is reported as a 7-day rolling average and color-coded by grouping common Pango lineage designations (https://cov-lineages.org) according to similar baseline Spike protein sequences, per the legend. Pango lineages are themselves evolving and represent diverse variants. These analyses done using the LANL COVID-19 Viral Genome Analysis Pipeline (www.cov.lanl.gov) using sequence data submitted to GISAID (Khare et al. China CDC Weekly 3(49):1049-1051 (2021)) and we used the the full GISAID data set (14,809,879 sequences) sampled globally between 2021/01/01 and 2023/05/10 to create this figure. Case count data was taken from the Johns Hopkins Center for Systems Science and Engineering (CSSEGIS (Dong et al. Lancet 20(5):533-534 (2020) https://github.com/CSSEGISandData/COVID-19), and then from Our World in Data (https://ourworldindata.org/covid-cases) when case count data began to become less reliable in early 2023 and so CSSEGIS stopped reporting it.

### A Spike mutations in VOCs

| WHO designation | Most common PANGO lineage | Most common mutational forms. NTD supersite, RBD, possibly furin cleavage related (positive charge near the furin cleavage site or H655Y), Heptad Repat 1 (HR1) |
|---|---|---|
| Alpha α | Alpha_B.1.1.7+Q.* | H69-,V70-,Y144-,N501Y,A570D,D614G,P681H,T716I,S982A,D1118H |
| Beta β | Beta_B.1.351 | D80A,D215G,L242-,A243-,L244-,K417N,E484K,N501Y,D614G,A701V |
| Gamma γ | Gamma_P.1 | L18F,T20N,P26S,D138Y,R190S,K417T,E484K,N501Y,D614G,H655Y,T1027I,V1176F |
| Delta δ | Delta_B.1.617.2+AY.* | T19R,T95I,G142D,E156-,F157-,R158G,L452R,T478K,D614G,P681R,D950N |
| Epsilon ε | B.1.429+B.1.427 | S13I,W152C,L452R,D614G |
| Iota ι | B.1.526 | L5F,T95I,D253G,E484K,D614G,A701V |
| Kappa κ | B.1.617.1 | T95I,G142D,E154K,L452R,E484Q,D614G,P681R,Q1071H |
| Lambda Λ | Lambda_C.37 | G75V,T76I,R246N,S247-,Y248-,L249-,T250-,P251-,G252-,D253-,L452Q,F490S,D614G,T859N |
| Mu μ | Mu_B.1.621 | T95I,+143T,Y144S,Y145N,R346K,E484K,N501Y,D614G,P681H,D950N |
| Omicron BA.1 o | Omicron B.1.1.529.1 | A67V,H69-,V70-,T95I,G142D,V143-,Y144-,Y145-, N211,L212I,+214EPE, G339D, S371L, S373P,S375F,K417N,N440K,G446S,S477N,T478K,E484A,Q493R,G496S,Q498R,N501Y, Y505H,T547K,D614G,H655Y,N679K,P681H,N764K,D796Y,N856K,Q954H,N969K,L981F |
| Omicron BA.2 | Omicron B.1.1.529.2 | T19I,L24-,P25-,P26-,A27S,G142D,V213G,G339D,S371F,S373P,S375F,T376A,D405N,R408S,K417N,N440K,S477N,T478K,E484A,Q493R,Q498R,N501Y,Y505H,D614G, H655Y,N679K,P681H,N764K,D796Y,Q954H,N969K |

### B Sites mutated in 2 or more VOCs

Redundant mutations

| Variant | E484 K Q A | L452 R Q | N501Y | K417 N T | T478K | T95I | G142D | Y144 del S | S371 F L | H69 V70 -> del69-70, V70F | H655Y | P681 R H | A701V | D950N |
|---|---|---|---|---|---|---|---|---|---|---|---|---|---|---|
| **D614G** | | | | | | | | | | | | | | |
| **Alpha** | | | N501Y | | | | | del144 | | del69-70 | | P681H | | |
| **Alpha_E484K** | E484K | | N501Y | | | | | del144 | | del69-70 | | P681H | | |
| **Delta_AY.3** | | L452R | | | T478K | T95I | G142D | | | | | **P681R** | | D950N |
| **AY.3_E484Q** | **E484Q** | L452R | | | T478K | T95I | G142D | | | | | **P681R** | | D950N |
| **AY.3_K417N** | | L452R | | K417N | T478K | T95I | G142D | | | | | **P681R** | | D950N |
| **Delta_AY.1** | | L452R | | K417N | T478K | T95I | G142D | | | | | **P681R** | | D950N |
| **Delta_AY.2** | | L452R | | K417N | T478K | T95I | G142D | | | **V70F** | | **P681R** | | D950N |
| **Beta** | E484K | | N501Y | **K417N** | | | | | | | | | A701V | |
| **Gamma** | E484K | | N501Y | **K417T** | | | | | | | H655Y | | | |
| **Iota** | E484K | | | | | T95I | | | | | | | A701V | |
| **Lambda** | | **L452Q** | | | | | | | | | | | | |
| **Epsilon** | | L452R | | | | | | | | | | | | |
| **Kappa** | **E484Q** | L452R | | | | T95I | G142D | | | | | **P681R** | | |
| **Mu** | E484K | | N501Y | | | T95I | | **Y144S** | | | | P681H | | D950N |
| **Omicron (BA.1)** | **E484A** | | N501Y | K417N | T478K | T95I | G142D | **del142-144** | S371L | del69-70 | H655Y | P681H | | |
| **BA.1+A484K** | E484K | | N501Y | K417N | T478K | T95I | G142D | **del142-144** | S371L | del69-70 | H655Y | P681H | | |
| **Omicron (BA.2)** | **E484A** | | N501Y | K417N | T478K | | G142D | | **S371F** | | H655Y | P681H | | |

**Red/Teal** = 2nd/3rd mutations; Mutation unique to single variant

### C Redundancy in mutations

Binary representation of mutations:

del69-70 = 1 if VOC has deletion at 69-70
0 if VOC does not have deletion at 69-70

del69-70 = del144 + del142-144
H655Y = K417T + S371L + S371F
P681H = del144 + Y144S + S371L + S371F
P681R = G142D – S371L – S37F
D950N = T478K + Y144S – S371L – S371F
A701V = N501Y - K417T + T95I - G142D - del144 - del142-144 - 2*Y144S

### D Spike mutations in holdout VOCs

| WHO designation | Most common PANGO lineage | Most common mutational forms. NTD supersite, RBD, possibly furin cleavage related (positive charge near the furin cleavage site or H655Y), Heptad Repat 1 (HR1) |
|---|---|---|
| BA.1.1 | Omicron B.1.1.529.1.1 | A67V,H69-,V70-,T95I,G142D,V143-,Y144-,Y145-,N211-,L212I,+214EPE,G339D,R346K, S371L,S373P,S375F,K417N,N440K,G446S,S477N,T478K,E484A,Q493R,G496S,Q498R, N501Y,Y505H,T547K,D614G,H655Y,N679K,P681H,N764K,D796Y,N856K,Q954H,N969K, L981F |
| BA.3 | Omicron B.1.1.529.3 | A67V,H69-,V70-,T95I,G142D,V143-,Y144-,Y145-,N211-,L212I,G339D,S371F,S373P, S375F,D405N,K417N,N440K,G446S,S477N,T478K,E484A,Q493R,Q498R,N501Y,Y505H, D614G,H655Y,N679K,P681H,N764K,D796Y,Q954H,N969K |
| BA.2.12.1 | Omicron B.1.1.529.2.12.1 | T19I,L24S,P25-,P26-,A27-,G142D,V213G,G339D,S371F,S373P,S375F,T376A,D405N, R408S,K417N,N440K,L452Q,S477N,T478K,E484A,Q493R,Q498R,N501Y,Y505H,D614G, H655Y,N679K,P681H,S704L,N764K,D796Y,Q954H,N969K |
| BA.4/BA.5 | Omicron B.1.1.529.4/5 | T19I,L24S,P25-,P26-,A27-,G142D,V213G,G339D,S371F,S373P,S375F,T376A,D405N, R408S,K417N,N440K,L452R,S477N,T478K,E484A,F486V,Q498R,N501Y,Y505H,D614G, H655Y,N679K,P681H,N764K,D796Y,Q954H,N969K |

**Figure S2: Spike sequence and site identification for models (related to Fig. 3, 5).** (A) Spike sequences of variant pseudoviruses used in the training dataset. The sequence for each infecting strain was assumed to be the same as that of the corresponding pseudovirus. (B) Mutations in each variant in our training dataset where at least 3 or more wildtype variants (i.e. not counting mutant pseudoviruses, e.g. Alpha + E484K) show mutations away from the prototype strain. Site 371 with mutations in BA.1 and BA.2 was added due to the strong resistance of S371L/F to most classes of RBD neutralizing antibodies. Sites with redundant mutations (see (C)) are indicated. (C) Linear dependence of mutations at sites 69-70, 655, 681, 701 and 950 as a function of mutations at 9 sites chosen for the model. (D) Same as (A), except Spike sequences for novel variants in holdout dataset are shown.

**A** Heterologous ID50 titer scatterplots

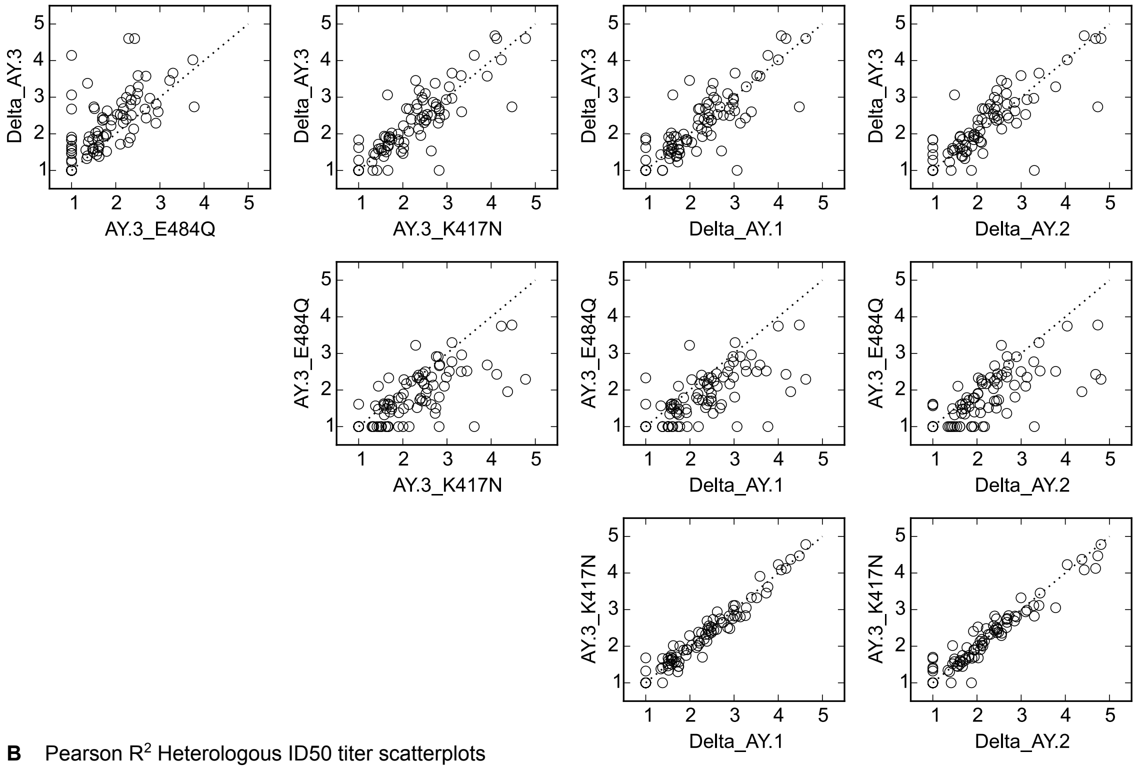


**B** Pearson $R^2$ Heterologous ID50 titer scatterplots

| | Delta_AY.3 | AY.3_E484Q | AY.3_K417N | Delta_AY.1 | Delta_AY.2 |
|---|---|---|---|---|---|
| Delta_AY.3 | 1.0000 | 0.5446 | 0.7759 | 0.7422 | 0.7488 |
| AY.3_E484Q | 0.5446 | 1.0000 | 0.6047 | 0.6039 | 0.6331 |
| AY.3_K417N | 0.7759 | 0.6047 | 1.0000 | 0.9560 | 0.9423 |
| Delta_AY.1 | 0.7422 | 0.6039 | 0.9560 | 1.0000 | 0.9115 |
| Delta_AY.2 | 0.7488 | 0.6331 | 0.9423 | 0.9115 | 1.0000 |

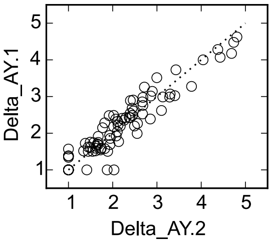


**Figure S3: Neutralization of Delta and its related variants/mutants (related to Fig. 1, 2).** (A) ID50 titers are shown for each pair of Delta-related pseudoviruses across all sera that were tested against each given pair of pseudoviruses. (B) Pearson $R^2$ values across pairs of pseudoviruses.

**A** Pseudovirus mutation model fits

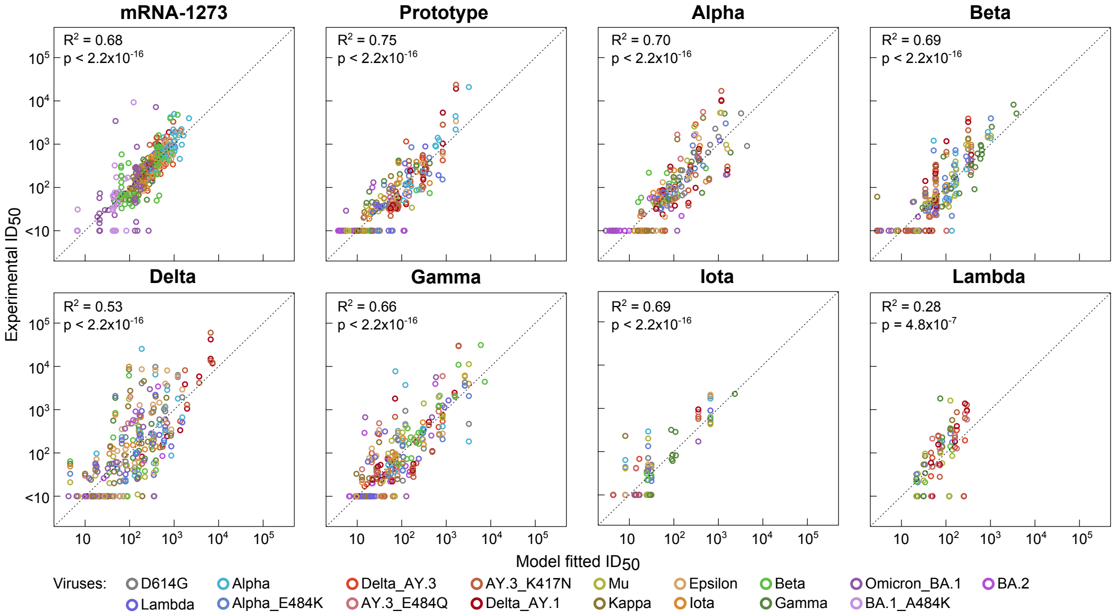


**B** Pseudovirus mutation model coefficients

| Sera model | Intercept | Auto ID50 | S371L | S371F | E484K | E484Q | E484A | L452R | L452Q | T478K | N501Y | K417N | K417T | T95I | G142D | del144 | Y144S |
|---|---|---|---|---|---|---|---|---|---|---|---|---|---|---|---|---|---|
| mRNA-1273 | **0.3397**<br>(0.0352) | **0.7792**<br>($<2.2\times10^{-16}$) | **-1.1824**<br>($<2.2\times10^{-16}$) | NA | **-0.4998**<br>($<2.2\times10^{-16}$) | **-0.4924**<br>($3.8\times10^{-12}$) | | | | | **0.1918**<br>(0.0032) | **-0.3311**<br>($4.2\times10^{-10}$) | | **0.1947**<br>(0.0126) | | | **-0.5166**<br>(0.0002) |
| Prototype | **0.4284**<br>(0.0003) | **0.6786**<br>($<2.2\times10^{-16}$) | **-1.1567**<br>($<2.2\times10^{-16}$) | **-1.1805**<br>($<2.2\times10^{-16}$) | **-0.5915**<br>($<2.2\times10^{-16}$) | **-0.7623**<br>($<2.2\times10^{-16}$) | NA | | **-0.3932**<br>(0.0002) | | | | | | | **0.2882**<br>(0.0002) | **-0.4007**<br>(0.0002) |
| Lambda | 0.4198 | **0.5749**<br>($7.2\times10^{-6}$) | NA | NA | | | NA | NA | NA | **0.3296**<br>(0.0124) | NA | | | | NA | | |
| Delta | 0.2241 | **0.5597**<br>($<2.2\times10^{-16}$) | **-1.6504**<br>($1.4\times10^{-15}$) | **-1.4658**<br>($1.4\times10^{-10}$) | **-0.5929**<br>($1.8\times10^{-10}$) | **-1.2781**<br>($1.4\times10^{-11}$) | NA | | | | | | | | **1.0199**<br>($4.6\times10^{-10}$) | | NA |
| Alpha | **0.3729**<br>(0.0377) | **0.7691**<br>($<2.2\times10^{-16}$) | **-1.5687**<br>($<2.2\times10^{-16}$) | **-1.4729**<br>($<2.2\times10^{-16}$) | **-0.9253**<br>($6.8\times10^{-10}$) | **-0.4108**<br>($6.0\times10^{-5}$) | NA | **-0.4421**<br>(0.0008) | **-0.4745**<br>(0.0041) | | | | **0.5548**<br>(0.0001) | | | **0.5806**<br>($9.9\times10^{-5}$) | **0.4551**<br>(0.0021) |
| Beta | **-0.4284**<br>($6.2\times10^{-5}$) | **0.8841**<br>($<2.2\times10^{-16}$) | **-0.8259**<br>($1.2\times10^{-12}$) | NA | | | NA | | | | **0.4421**<br>($5.2\times10^{-11}$) | | **0.5765**<br>($3.7\times10^{-7}$) | | | | NA |
| Gamma | 0.0256 | **0.7604**<br>($<2.2\times10^{-16}$) | **-1.0967**<br>($2.2\times10^{-12}$) | **-1.1125**<br>($5.9\times10^{-12}$) | | | NA | **-0.5897**<br>($1.3\times10^{-12}$) | **-0.4576**<br>(0.0006) | | | **0.3586**<br>($2.0\times10^{-5}$) | NA | | | | |
| Iota | 0.0250 | **0.8850**<br>($<2.2\times10^{-16}$) | | NA | | | NA | | | **-0.2719**<br>(0.0012) | | | **0.5491**<br>(0.0003) | | | | |

**C** Crossvalidation prediction accuracy $R^2$ for mismatch and pseudovirus mutation models

| CV | Model | mRNA-1274 | Prototype | Alpha | Beta | Gamma | Delta | Iota | Lambda | Average across inf./vac. strain |
|---|---|---|---|---|---|---|---|---|---|---|
| Virus CV | Mismatch BIC | 0.4374 | 0.4980 | 0.5078 | 0.5409 | 0.5857 | 0.3202 | 0.4149 | 0.2329 | 0.4422 |
| | Mutations BIC | 0.5222 | 0.1671 | 0.0000 | 0.5409 | 0.3923 | 0.0000 | 0.6182 | 0.2538 | 0.3118 |
| Serum CV | Mismatch BIC | 0.6174 | 0.6682 | 0.5238 | 0.6209 | 0.6039 | 0.4414 | 0.0000 | 0.0000 | 0.4345 |
| | Mutations BIC | 0.6440 | 0.7080 | 0.5623 | 0.6209 | 0.6046 | 0.4675 | 0.0000 | 0.0000 | 0.4509 |
| Average Virus + Serum CV | Mismatch BIC | 0.5274 | 0.5831 | 0.5158 | 0.5809 | 0.5948 | 0.3808 | 0.2074 | 0.1165 | 0.4383 |
| | Mutations BIC | 0.5831 | 0.4375 | 0.2812 | 0.5809 | 0.4985 | 0.2338 | 0.3091 | 0.1269 | 0.3814 |

**Figure S4: Infecting or vaccine strain specific models using mutations in pseudoviruses (related to Fig. 3).** (A) Experimental ID50 titers (vertical axis) are plotted against model fitted values (horizontal axis) for each infecting/vaccine strain. Each model was selected using backward model selection Coefficient of determination $R^2$ and p-values from Kendall tau rank test are shown in each sub-panel. Different colors indicate different pseudoviruses per the legend. (B) Coefficients for each variable (columns) are shown for each infecting/vaccine strain specific model (rows), together with the Student's t-test p-values in parentheses below for significant ($p < 0.05$) variables. Coefficients are color-coded using the red-white-blue color scale corresponding to negative, 0 and positive values. A blank entry indicates a variable that was dropped using the BIC model reduction strategy. “NA” indicates a variable that either had no variation or was redundant (i.e. linearly explained by other variables) for the data available for that infecting/vaccine strain. (C) Comparison of predictive accuracy of “mismatch models” (Fig. 3) to the mutation models in this figure using pseudovirus crossvalidation (CV) and leave-one-out CV on each serum for a given infecting/vaccine strain. The table entries show coefficient of determination ($R^2$) values color-coded using a red-white-blue scheme that corresponds to 0, 0.5 and 1.0, respectively. The last column indicates average over all infecting/vaccine strain models, and the bottom two rows indicate average across both pseudovirus and serum CV. The rightmost entries in the bottom two rows indicate average across both types of CV as well as across infecting/vaccine strains.

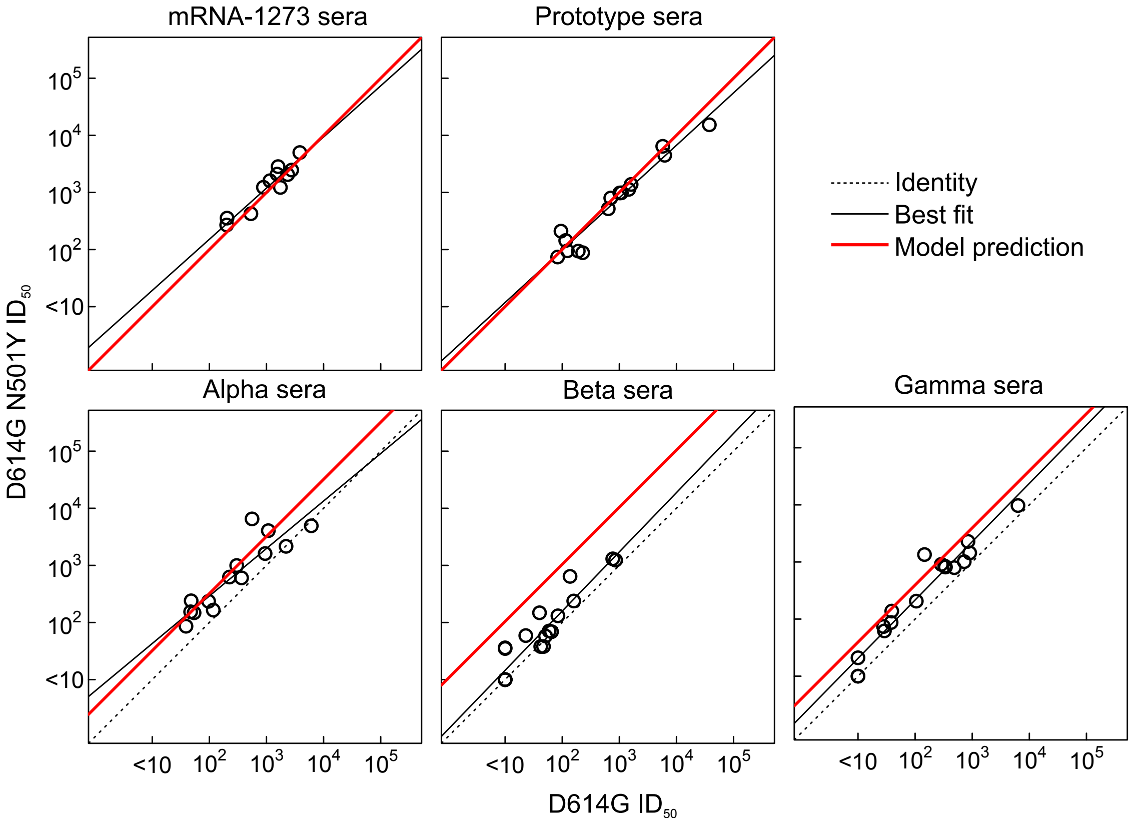


**Figure S5: Impact of N501Y on neutralization by mRNA-1273 sera and by convalescent sera from Prototype, Alpha, Beta and Gamma infections (related to Fig. 3).** ID50 titers against D614G are shown on the horizontal axis and those against D614G+N501Y on the vertical axis. Dotted lines indicates identical titers for the two pseudoviruses, solid black lines indicate the best fit linear regression trend and red lines indicate the corresponding model predictions based on the models from Fig. 3C.

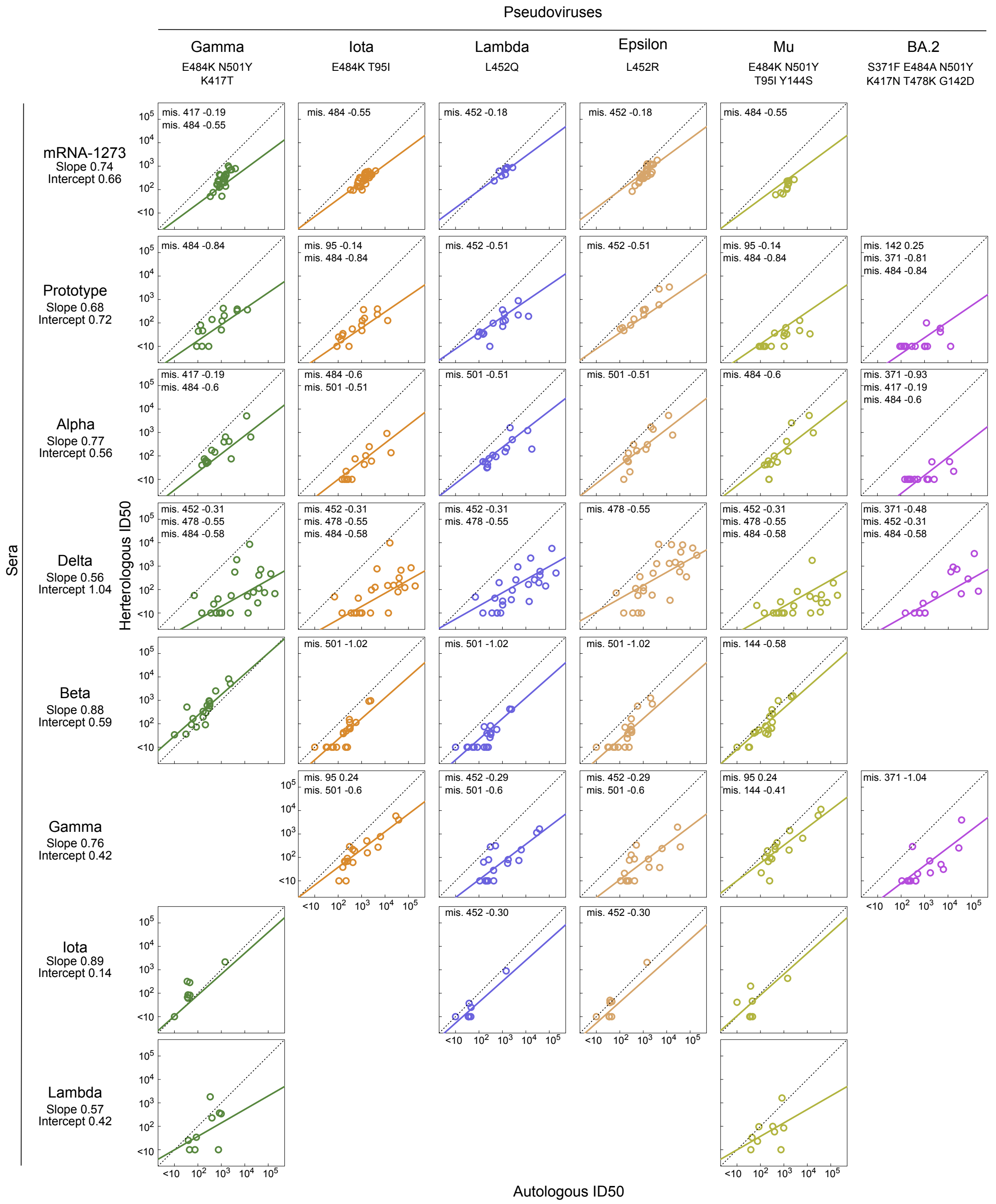


**Figure S6: Neutralization titers for sera from each infecting/vaccine strain (rows) against pseudoviruses (columns) with model fits from infecting/vaccine variant specific models (related to Fig. 4).** Same as Fig. 4, but for pseudoviruses not included in Fig. 4.

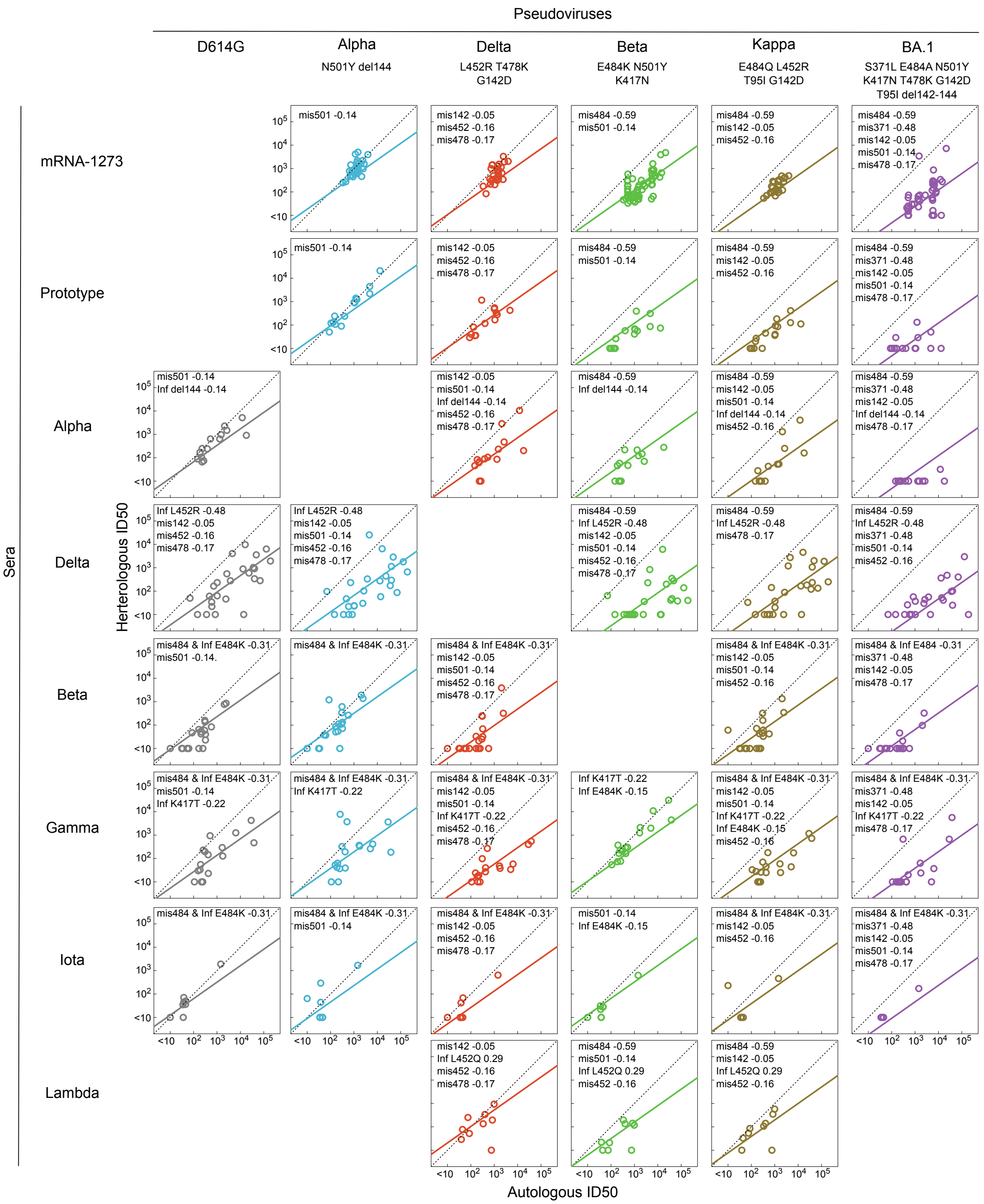

Pseudoviruses
D614G
Alpha
N501Y del144
Delta
L452R T478K
G142D
Beta
E484K N501Y
K417N
Kappa
E484Q L452R
T95I G142D
BA.1
S371L E484A N501Y
K417N T478K G142D
T95I del142-144
Sera
mRNA-1273
Prototype
Alpha
Delta
Beta
Gamma
Iota
Lambda
Herterologous ID50
Autologous ID50
mis501 -0.14
mis142 -0.05
mis452 -0.16
mis478 -0.17
mis484 -0.59
Inf del144 -0.14
mis371 -0.48
Inf L452R -0.48
mis484 & Inf E484K -0.31
mis484 & Inf E484 -0.31
Inf K417T -0.22
Inf E484K -0.15
Inf L452Q 0.29

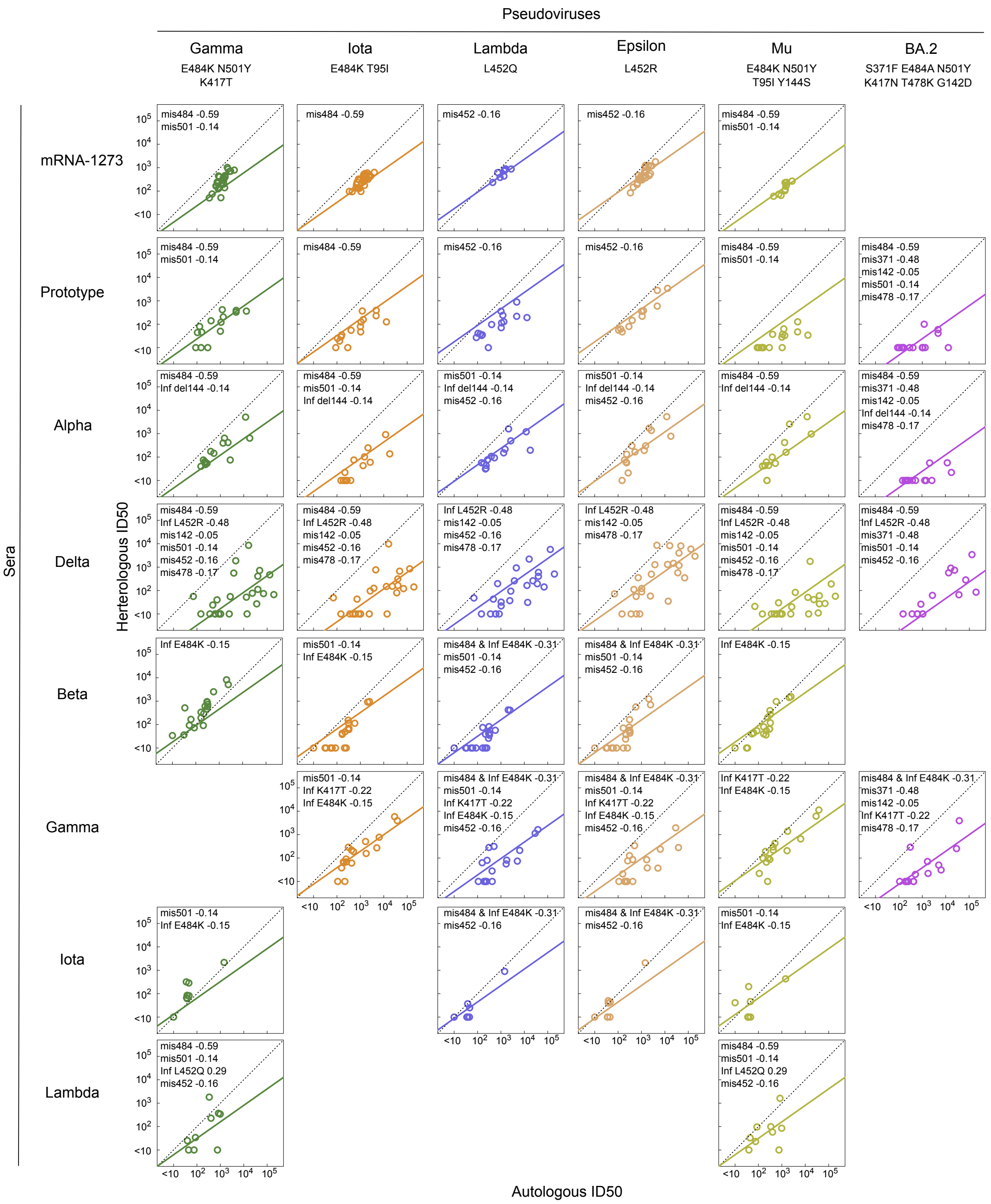


**Figures S7 & S8: Neutralization titers for sera from each infecting/vaccine strain (columns) against pseudoviruses (rows).** Same as Fig. 4, except the unified model from Fig. 5 is used to show the model fit lines and the mismatch and infecting strain variables that operate for a given infecting/vaccine strain and pseudovirus strain are shown with their coefficients.

**Text S1 – Model formulae and summaries**

**Formulae:**
All formulae below are in R's formula formats, where '~' is '=', '+' indicates linear combination, and '*' indicates pairwise-interaction terms that also add the individual terms. The coefficients for each variable that are fitted are implicitly assumed, and are not shown. Key for variable names: "Logid" = heterologous log10 ID50 titer; "Autoid" = autologous log10 ID50 titer; "mis_xxx" = binary variable encoding mismatch between infecting/vaccine strain and pseudovirus strain at position "xxx" (1=mismatch, 0=no mismatch); "Ser_X.YYY" = binary variable encoding whether the infecting/vaccine strain has the mutation "X" at site "YYY" (1=mutation, 0=no mutation or other than "X").

Autologous ID50 titer only model: Logid ~ Autoid

Mismatch only model: Logid ~ Autoid + mis_371 + mis_484 + mis_452 + mis_95 + mis_501+mis_142 + mis_144 + mis_417 + mis_478

Mismatch + Infecting strain model:
Logid ~ Autoid + mis_371 + mis_484 + mis_452 + mis_95 + mis_501+mis_142 + mis_144 + mis_417 + mis_478 + Ser_L.371 + Ser_K.484 + Ser_Q.484 + Ser_Y.501 + Ser_R.452 + Ser_Q.452 + Ser_I.95 + Ser_N.417 + Ser_T.417 + Ser_d.144 + Ser_S.144 + Ser_D.142 + Ser_K.478

Mismatch + infecting strain + interactions at same site model:
Logid ~ Autoid + mis_371*(Ser_L.371 ) + mis_484*(Ser_K.484 + Ser_Q.484) + mis_452*(Ser_R.452 + Ser_Q.452) + mis_95*(Ser_I.95) + mis_501*Ser_Y.501 + mis_142*Ser_D.142 + mis_144*(Ser_d.144 + Ser_S.144) + mis_417*(Ser_N.417+ Ser_T.417) + mis_478*Ser_K.478

Mismatch + infecting strain + interactions among all sites:
Logid ~ Autoid +
( Ser_L.371+Ser_K.484 + Ser_Q.484 + Ser_Y.501 + Ser_R.452 + Ser_Q.452 + Ser_I.95 + Ser_N.417 + Ser_T.417 + Ser_d.144 + Ser_S.144 + Ser_D.142 + Ser_K.478) *
(mis_484 + mis_452 + mis_95 + mis_501+mis_142 + mis_144 + mis_417 + mis_478)

Pseudovirus mutation model:
Logid ~ Autoid + Test_L.371 + Test_F.371 + Test_K.484 + Test_Q.484 + Test_A.484 + Test_R.452 + Test_Q.452 + Test_Y.501 + Test_N.417 + Test_T.417 + Test_K.478 + Test_I.95 + Test_D.142 + Test_d.144 + Test_S.144)

**Model summaries:**

Model summaries from R's summary function are shown below for each model from Figures 3 and 5. Terms like 'var1:var2' below indicate the interaction term between 'var1' and 'var2' that in our case of binary variables is just equal to the logical and operation (i.e. var1:var2 = 1 only if var1=1 and var2=1, and var1:var2 = 0 in all other cases).

mRNA-1273 BIC reduced model:

```
Deviance Residuals:
     Min        1Q    Median        3Q       Max
-1.18205  -0.17354  -0.01712   0.19797   2.05493

Coefficients:
            Estimate Std. Error t value Pr(>|t|)
(Intercept)  0.66211    0.17275   3.833 0.000151 ***
Autoid       0.73723    0.05304  13.899  < 2e-16 ***
mis_371     -0.78086    0.05842 -13.366  < 2e-16 ***
mis_484     -0.55141    0.05215 -10.574  < 2e-16 ***
mis_452     -0.17823    0.05258  -3.390 0.000781 ***
mis_417     -0.18767    0.04919  -3.815 0.000161 ***
---
Signif. codes:
0 ‘***’ 0.001 ‘**’ 0.01 ‘*’ 0.05 ‘.’ 0.1 ‘ ’ 1

(Dispersion parameter for gaussian family taken to be 0.1307885)

    Null deviance: 125.720  on 350  degrees of freedom
Residual deviance:  45.122  on 345  degrees of freedom
AIC: 290.05
```

Prototype sera BIC reduced model:

```
Deviance Residuals:
     Min        1Q    Median        3Q       Max
-1.12646  -0.24089  -0.02086   0.23799   1.15768

Coefficients:
            Estimate Std. Error t value Pr(>|t|)
(Intercept)  0.72121    0.13522   5.334 2.48e-07 ***
Autoid       0.68219    0.04029  16.930  < 2e-16 ***
mis_371     -0.80973    0.11967  -6.766 1.29e-10 ***
mis_484     -0.84309    0.07060 -11.942  < 2e-16 ***
mis_452     -0.51463    0.10566  -4.871 2.19e-06 ***
mis_95      -0.14080    0.06125  -2.299   0.0225 *
mis_142      0.25151    0.09201   2.733   0.0068 **
---
Signif. codes:
0 ‘***’ 0.001 ‘**’ 0.01 ‘*’ 0.05 ‘.’ 0.1 ‘ ’ 1

(Dispersion parameter for gaussian family taken to be 0.146332)

    Null deviance: 110.55  on 216  degrees of freedom
Residual deviance:  30.73  on 210  degrees of freedom
AIC: 207.66
```

Alpha sera BIC reduced model:

```
Deviance Residuals:
     Min        1Q    Median        3Q       Max
-1.13006  -0.27843  -0.02328   0.31034   1.21443

Coefficients:
            Estimate Std. Error t value Pr(>|t|)
(Intercept)  0.55685    0.18389   3.028  0.00278 **
Autoid       0.77312    0.04808  16.081  < 2e-16 ***
mis_371     -0.93038    0.11104  -8.378 9.19e-15 ***
```

```
mis_484     -0.59867    0.08818  -6.789 1.24e-10 ***
mis_501     -0.50711    0.10045  -5.049 9.95e-07 ***
mis_417     -0.18753    0.07885  -2.378  0.01832 *
---
Signif. codes:
0 ‘***’ 0.001 ‘**’ 0.01 ‘*’ 0.05 ‘.’ 0.1 ‘ ’ 1

(Dispersion parameter for gaussian family taken to be 0.1983889)

    Null deviance: 123.080  on 206  degrees of freedom
Residual deviance:  39.876  on 201  degrees of freedom
AIC: 260.52

Number of Fisher Scoring iterations: 2
```

Beta sera BIC reduced model:

```
Deviance Residuals:
     Min        1Q    Median        3Q       Max
-1.13220  -0.28193  -0.02706   0.22387   1.35590

Coefficients:
            Estimate Std. Error t value Pr(>|t|)
(Intercept)  0.59021    0.13831   4.267 2.78e-05 ***
Autoid       0.88412    0.04442  19.902  < 2e-16 ***
mis_371     -0.82591    0.11049  -7.475 1.21e-12 ***
mis_501     -1.01860    0.10047 -10.139  < 2e-16 ***
mis_144     -0.57653    0.11049  -5.218 3.73e-07 ***
---
Signif. codes:
0 ‘***’ 0.001 ‘**’ 0.01 ‘*’ 0.05 ‘.’ 0.1 ‘ ’ 1

(Dispersion parameter for gaussian family taken to be 0.1723962)

    Null deviance: 140.713  on 261  degrees of freedom
Residual deviance:  44.306  on 257  degrees of freedom
AIC: 289.89
```

Gamma sera BIC reduced model:

```
Deviance Residuals:
    Min       1Q   Median       3Q      Max
-1.2126  -0.3110  -0.0574   0.2322   2.0548

Coefficients:
            Estimate Std. Error t value Pr(>|t|)
(Intercept)  0.41871    0.15973   2.621 0.009317 **
Autoid       0.75830    0.04064  18.661  < 2e-16 ***
mis_371     -1.04027    0.11539  -9.015  < 2e-16 ***
mis_452     -0.29311    0.10222  -2.868 0.004505 **
mis_95       0.23872    0.08320   2.869 0.004482 **
mis_501     -0.59571    0.14748  -4.039 7.22e-05 ***
mis_144     -0.41231    0.12104  -3.407 0.000771 ***
---
Signif. codes:
0 ‘***’ 0.001 ‘**’ 0.01 ‘*’ 0.05 ‘.’ 0.1 ‘ ’ 1

(Dispersion parameter for gaussian family taken to be 0.2448321)
```

```
    Null deviance: 170.69  on 246  degrees of freedom
Residual deviance:  58.76  on 240  degrees of freedom
AIC: 362.28
```

Delta sera BIC reduced model:

```
Deviance Residuals:
     Min        1Q    Median        3Q       Max
-1.78009  -0.40352  -0.09826   0.40187   2.17147

Coefficients:
            Estimate Std. Error t value Pr(>|t|)
(Intercept)  1.03570    0.18735   5.528 7.14e-08 ***
Autoid       0.56196    0.03964  14.175  < 2e-16 ***
mis_371     -0.48372    0.19556  -2.473 0.013946 *
mis_484     -0.58376    0.08178  -7.138 7.43e-12 ***
mis_452     -0.31159    0.10361  -3.007 0.002863 **
mis_478     -0.54628    0.15188  -3.597 0.000378 ***
---
Signif. codes:
0 '***' 0.001 '**' 0.01 '*' 0.05 '.' 0.1 ' ' 1

(Dispersion parameter for gaussian family taken to be 0.4187832)

    Null deviance: 260.00  on 299  degrees of freedom
Residual deviance: 123.12  on 294  degrees of freedom
AIC: 598.18
```

Iota sera BIC reduced model:

```
Deviance Residuals:
    Min       1Q   Median       3Q      Max
-0.6198  -0.2439  -0.1228   0.2736   1.6372

Coefficients:
            Estimate Std. Error t value Pr(>|t|)
(Intercept)  0.13574    0.13019   1.043 0.299616
Autoid       0.89251    0.06754  13.215  < 2e-16 ***
mis_371     -0.59787    0.21416  -2.792 0.006272 **
mis_452     -0.30183    0.08171  -3.694 0.000358 ***
---
Signif. codes:
0 '***' 0.001 '**' 0.01 '*' 0.05 '.' 0.1 ' ' 1

(Dispersion parameter for gaussian family taken to be 0.1681447)

    Null deviance: 48.713  on 104  degrees of freedom
Residual deviance: 16.983  on 101  degrees of freedom
AIC: 116.69
```

Lambda sera BIC reduced model:

```
Deviance Residuals:
    Min       1Q   Median       3Q      Max
-1.4033  -0.2097   0.1234   0.3725   1.3855

Coefficients:
            Estimate Std. Error t value Pr(>|t|)
```

```
(Intercept)   0.4198     0.2912   1.441   0.1535
Autoid        0.5749     0.1195   4.812 7.17e-06 ***
mis_478       0.3296     0.1287   2.560   0.0124 *
---
Signif. codes:
0 '***' 0.001 '**' 0.01 '*' 0.05 '.' 0.1 ' ' 1

(Dispersion parameter for gaussian family taken to be 0.3315181)

    Null deviance: 35.707  on 80  degrees of freedom
Residual deviance: 25.858  on 78  degrees of freedom
AIC: 145.38
```

Unified mismatch + infecting strain + interactions at same site BIC forward model:

```
Deviance Residuals:
     Min        1Q    Median        3Q       Max
-1.76476  -0.30053  -0.02355   0.28298   2.12625

Coefficients:
                  Estimate Std. Error t value Pr(>|t|)
(Intercept)        0.70237    0.06371  11.024  < 2e-16 ***
Autoid             0.70379    0.01747  40.275  < 2e-16 ***
mis_484           -0.59299    0.03460 -17.141  < 2e-16 ***
Ser_R.452         -0.47539    0.04051 -11.736  < 2e-16 ***
mis_371           -0.47695    0.04362 -10.934  < 2e-16 ***
mis_142           -0.05144    0.05300  -0.971 0.331877
mis_501           -0.14472    0.02539  -5.700 1.40e-08 ***
Ser_d.144         -0.13821    0.04099  -3.371 0.000764 ***
Ser_T.417         -0.22370    0.04371  -5.118 3.43e-07 ***
Ser_K.484         -0.15319    0.05697  -2.689 0.007232 **
Ser_Q.452          0.28535    0.06367   4.481 7.89e-06 ***
mis_452           -0.15884    0.03489  -4.552 5.67e-06 ***
mis_478           -0.16766    0.05069  -3.307 0.000961 ***
mis_484:Ser_K.484  0.43899    0.06377   6.884 8.06e-12 ***
---
Signif. codes:
0 '***' 0.001 '**' 0.01 '*' 0.05 '.' 0.1 ' ' 1

(Dispersion parameter for gaussian family taken to be 0.2519931)

    Null deviance: 1105.4  on 1769  degrees of freedom
Residual deviance:  442.5  on 1756  degrees of freedom
AIC: 2599.3
```
